# A multi-omics analysis of human fibroblasts overexpressing an *Alu* transposon reveals widespread disruptions in aging-associated pathways

**DOI:** 10.1101/2025.07.11.664466

**Authors:** Juan I. Bravo, Eyael Tewelde, Christina D. King, Joanna Bons, Samah Shah, Jacob Rose, Judith Campisi, Birgit Schilling, Bérénice A. Benayoun

## Abstract

During aging and cellular senescence, repetitive elements are frequently transcriptionally derepressed across species and cell types. Among these, the most abundant repeats by copy number in the human genome are *Alu* retrotransposons. Though *Alu* elements are often studied for their mutagenic potential, there is increasing appreciation for their contributions to other biological functions, including pro-inflammatory signaling and mitochondrial dysfunction. However, a comprehensive analysis of *Alu*-driven molecular changes remains to be conducted, and *Alu*’s potential contributions to aging features remain incompletely characterized. Here, we show that overexpression of an *AluJb* transposon in human primary IMR-90 fibroblasts leads to large-scale alterations across the transcriptome, cellular proteome, and secretome. Functional genomics analyses reveal alterations in aging pathways, broadly, and mitochondrial metabolism, proteostasis, cell cycle, and extracellular matrix pathways, more specifically. Our results demonstrate that *Alu* transcriptional upregulation is sufficient to drive widespread disruptions to cellular homeostasis that mirror aging-associated alterations.

## INTRODUCTION

Transposable elements (TEs), often informally referred to as “jumping genes”, are genetic elements that can mobilize within a genome. Depending on mobilization mechanisms, TEs can be classified as DNA transposons, which operate through a “cut-and-paste” mechanism, or retrotransposons, which operate through a “copy-and-paste” mechanism [1]. Historically, transposable elements have been largely ignored in biological studies because of (1) biases that considered TEs “junk” DNA and (2) the extreme difficulty of working with multi-copy, highly repetitive genomic sequences. Thus, there is frequently a gap in our understanding of the relationship between TEs and biological processes of interest, such as cellular senescence or aging.

Technological advancements over the past two decades (including next generation sequencing) have facilitated the study of TEs, including in the context of aging and aging-associated diseases [reviewed in [2]]. For example, age-related de-repression of TEs has been observed across diverse model systems, including *Caenorhabditis elegans* [3], *Drosophila* [4–6], and *Mus musculus* [7, 8], and in senescent cells [9–11]. Importantly, TE de-repression has also been observed across age-associated pathologies, e.g. cell senescence, cancer [12–15], and Alzheimer’s disease [16, 17]. These findings suggest that aging biology and transposon biology are tightly intertwined. However, whether TEs are drivers of aging and aging phenotypes or whether they are mere “innocent” bystanders of the aging process is still unknown.

In the human genome, the two most abundant families of TEs are the retrotransposons long interspersed element 1 (*LINE-1* or *L1*) and *Alu*. *Alu* transposons, in particular, have the largest number of copies, with ∼1.1 million loci occupying ∼9-11% of the genome [18, 19]. These retrotransposons are short (∼300 base pairs) and require the *LINE-1* machinery for their own mobilization [20–22]. Further, the *Alu* family can be segregated into three subfamilies depending on evolutionary age: *AluJ* is the oldest lineage and likely immobile in humans, *AluS* is the middle-aged lineage containing some mobile copies, and *AluY* is the youngest lineage with the most functionally intact copies [23]. Evolutionarily, *Alu* elements are believed to have originated from 7SL RNA [20, 24] which plays important roles in (1) co-translationally directing proteins to the endoplasmic reticulum as part of the protein secretory pathway [25–27] and (2) orchestrating a cellular response to thermal stress that involves transcription and translation arrest [28]. Its evolutionary history suggests that *Alu* may play important roles in regulating proteostasis, although this has not been thoroughly evaluated, and the breadth of *Alu*-regulated pathways remains unclearly defined.

Thus far, the effects of elevated *Alu* expression in a non-diseased and non-senescent background are ill-defined, and whether *Alu* overexpression is sufficient to recapitulate aging phenotypes is an open question. To address these gaps, we generated a multi-omics resource encompassing transcriptomic, proteomic, and secretomic profiles of “healthy” human primary IMR-90 fibroblasts overexpressing an *AluJb* retrotransposon, an element which naturally becomes transcriptionally upregulated in human primary aging fibroblasts. We demonstrate that this overexpression induces widescale remodeling of the transcriptome, cellular proteome, and secretome, and that this remodeling partially reflects remodeling observed during aging. On both a molecular and functional level, we observe *AluJb*-induced alterations in mitochondrial metabolic activity, cell proliferation, and proteostasis—all pathways that become mis-regulated during aging. Our results suggest that *AluJb* overexpression is sufficient to regulate the expression and functional output of aging-associated pathways, and, therefore, *AluJb* misregulation with aging may contribute to the onset of cellular aging phenotypes when de-repressed under physiological conditions with aging.

## MATERIAL AND METHODS

### Cell lines and cell culture conditions

IMR-90 (Coriell cat. I90-83, RRID: CVCL_0347) human female embryonic fibroblast cells were sourced from the Coriell Institute. Cells were maintained in Minimum Essential Medium (MEM) containing Earle’s salts (Corning cat. 15-010-CV), 15% fetal bovine serum (FBS, Sigma-Aldrich cat. F0926-500ML), 1X non-essential amino acids (NEAA, Quality Biological cat. 116-078-721), and 1X Penicillin-Streptomycin-Glutamine (Corning cat. 30-009-CI). Cells were cultured in a humidified incubator at 37°C and 5% CO_2_, subculturing cells 1:6 once they reached ∼90% confluency. All cells used were maintained below passage 30. Specifically, multi-omics samples on quiescent cells were prepared from cells at passage 13, and transcriptomic analyses with proliferating cells were carried out at passage 10. Cells were routinely tested for mycoplasma contamination using the PlasmoTest Mycoplasma Detection Kit (InvivoGen).

The identity and purity of cells used in this study were verified using ATCC’s Human Cell STR Profiling Service (ATCC 135-XV).

### Plasmid Construction

The empty pLKO.1 cloning vector, which contains the U6 promoter, was a gift from David Root (Addgene plasmid #10878; http://n2t.net/addgene:10878; RRID: Addgene_10878). The empty pUC19 cloning vector was purchased from New England Biolabs (NEB; cat. N3041S).

The *AluJb* overexpression construct was generated by overlap extension polymerase chain reaction (PCR). To begin, the *AluJb* consensus sequence was obtained from Repbase [29] and submitted to Genewiz for gene synthesis, yielding pUC57_AluJb. Next, the U6 promoter was amplified from pLKO.1 using primers TAAGCA_HindIII_U6_For and AluJB_exten_U6_R, and *AluJb* was amplified from pUC57_AluJb using primers U6_exten_AluJB_F and TAAGCA_EcoRI_5T_AluJB_R. Importantly, the *AluJb* amplification primers introduced a stretch of 5 thymidine residues in the *AluJb* 3’ terminal region to act as a minimal RNA polymerase III termination signal [30]. A first round of PCRs was carried out using AccuPrime *Pfx* DNA polymerase (Invitrogen cat. 12344024) in a Biorad C1000 Touch Thermal Cycler with these steps: initial denaturation (95°C, 2 minutes) and 35 cycles of denaturation (95°C, 15 seconds), annealing (55°C, 30 seconds), and extension (68°C, 60 seconds). The PCR products were run on an agarose gel, the amplicons of the expected size were excised, and the DNA was purified using the Nucleospin Gel and PCR Cleanup Kit (Macherey-Nagel item 740609.50). After, equimolar amounts of the U6 amplicon and the *AluJb* amplicon were mixed together with *Pfx* DNA polymerase, in order to fuse the U6 promoter with *AluJb* through a second round of PCRs (initial denaturation at 98°C for 30 seconds and 7 cycles of denaturation at 98°C for 10 second, annealing at 72°C for 30 second, and extension at 72°C for 15 seconds). An aliquot of the resulting PCR products was used as a template for a third round of PCRs to amplify the final 541 base pair U6_AluJb fusion product using primers TAAGCA_HindIII_U6_For and TAAGCA_EcoRI_5T_AluJB_R. This third set of PCRs was run under the same conditions as the first round but only for 25 cycles and using an annealing temperature of 60°C. Again, PCR products were run on an agarose gel and purified as before. Then, the pUC19 backbone and purified U6_AluJb amplicon were both digested with HindIII-HF and EcoRI-HF, the desired digestion products were purified from an agarose gel, and the backbone and insert were ligated together using the Quick Ligation Kit (NEB cat. M2200S), generating pUC19-U6_AluJb. NEB Turbo competent *Escherichia coli* (*E. coli*) (cat. C2984H) were transformed with the ligation product using the manufacturer recommended high efficiency transformation protocol. The final U6_AluJb fusion on the pUC19 backbone was sequence verified using Sanger sequencing. For a control plasmid, we introduced only the U6 promoter onto the pUC19 backbone (pUC19-U6) following the protocol outlined above with these modifications: (i) the U6 promoter was amplified with primers TAAGCA_HindIII_U6_For and TAAGCA_EcoRI_U6_R during the first PCRs and then gel-purified, and (ii) the second and third PCRs were skipped since the promoter was not fused with any other amplicon. The U6 promoter insert on the pUC19 backbone was also sequence verified using Sanger sequencing.

The plasmids generated in this study, pUC19-U6 (ID: 240313) and pUC19-U6_AluJb (ID: 240314), are available through Addgene.

### Transfections and sample collections for ‘omics’ analyses

*E. coli* were cultured in LB Broth (Thermo Fisher Scientific) supplemented with 50 μg/mL carbenicillin to an optical density 600 (OD_600_) of 2 – 4. Plasmid extractions were carried out using the Nucleobond Xtra Midi Plus EF kit (Macherey-Nagel item no. 740422.50) following manufacturer recommendations. Plasmids were aliquoted and stored at −20°C until the time of transfection.

On the day of transfection, we washed IMR-90 fibroblasts twice with Dulbecco’s phosphate-buffered saline (DPBS, Corning cat. 21-031-CV), detached cells using 0.25% trypsin (HyClone cat. 95053-258), neutralized the trypsin using a two-fold higher volume of media, spun cells down (500xG, 5 minutes), resuspended cells in fresh media, and counted cells by trypan blue staining using a Countess II FL automated cell counter (Thermo Fisher) or a CellDrop Automated Cell Counter (DeNovix). The number of cells necessary for the experiment were then aliquoted, spun down, and washed once with DPBS. Fibroblasts were transfected by electroporation using the Neon Transfection System (Invitrogen) with the following parameters: Buffer R for cell resuspension, 1500 V, 30 ms, and 1 pulse. Per reaction, we maintained a plasmid mass : cell number ratio of 2.5 μg : 2*10^6^ cells. We independently transfected 1.0 – 1.2 * 10^7^ fibroblasts per 100 mm tissue culture-treated petri dish (GenClone cat. 25-202), preparing 1 – 2 dishes to be used for each biological replicate downstream. Immediately after transfection, cells were cultured in Penicillin-Streptomycin-free media and allowed to recover from the electroporation for ∼24 hours. Since cell viability following electroporation exhibited plate-to-plate variability, surviving cells were then detached by trypsinization, counted, and re-seeded at equal densities across biological replicates and experimental conditions. We subsequently defined this time point as “0 hours”.

For paired transcriptomic and proteomic analyses, fibroblasts were prepared following previously described guidelines for generating conditioned media from senescent cells and quiescent control cells [31]. We implemented this protocol because (i) we hypothesized that *Alu* overexpression may induce senescence-associated changes, based on the published literature, and (ii) this protocol utilizes serum deprivation to restrict proliferation in order to isolate senescent versus non-senescent effects by eliminating non-proliferating vs proliferating effects. More specifically, fibroblasts were independently re-seeded at a density of 6 * 10^6^ cells per 150 mm tissue culture-treated dish (Corning cat. 430599), with 5 biological replicates per experimental group. Cells were cultured in 60 mL of 0.2% FBS media, which was replaced after 24 hours. At 48 hours, the media was removed, cells were washed twice with DPBS, and cells were fed with 60 mL of FBS-free and phenol red-free MEM media (Corning cat. 17-305-CV) containing Penicillin-Streptomycin-Glutamine and non-essential amino acids. At 72 hours, supernatants were collected in conicals, conicals were spun down (4000xG, 5 minutes, 4°C) to pellet cell debris, and supernatants were processed through Target2 0.45 μm polyvinylidene fluoride (PVDF) syringe filters (Thermo Scientific cat. F2500-5) to remove any remaining cellular debris. Filtered conditioned media was snap frozen in liquid nitrogen and stored at −80°C. Cells were washed with DPBS and detached with 0.05% trypsin (Corning cat. 25-052-CI), trypsin was neutralized with FBS-containing media, and cells were spun down. Cells were washed once with FBS-free and phenol red-free media, spun down, and resuspended again in that same media. Cell viability was assessed by trypan blue staining, ensuring that cell viability > 90% for all samples, and each sample was aliquoted into two tubes, saving 40% of the cells for proteomic analysis and 60% for transcriptomic analysis. Cells were spun down, the supernatants were discarded, and the cell pellets were snap frozen in liquid nitrogen and stored at −80°C. Cellular pellets for transcriptomic analysis were later lysed in TRIzol Reagent (Invitrogen) for downstream total RNA isolation (see below).

To ensure that major *Alu*-induced pathway changes replicated in proliferating cells, we prepared additional transcriptomic samples for IMR-90 fibroblasts overexpressing *AluJb* in standard, serum-containing maintenance media. More specifically, fibroblasts were independently transfected as before. Following recovery from the electroporation, fibroblasts were independently re-seeded at a density of 2 – 3 * 10^6^ fibroblasts per 100 mm dish containing maintenance media, with 4 biological replicates per experimental group. After 24 hours, cells were lysed in TRIzol Reagent (Invitrogen) for downstream total RNA isolation (see below).

### RNA extractions and mRNA sequencing

RNA was extracted using the Direct-zol RNA Miniprep kit (Zymo Research cat. R2052) following manufacturer recommendations. To confirm *AluJb* overexpression, complementary DNA (cDNA) was generated for each RNA sample using the Maxima H Minus cDNA Synthesis Master Mix with dsDNase (Thermo Scientific cat. M1682), following manufacturer instructions and including reverse transcriptase negative (RT-) controls. We verified plasmid-specific *AluJb* overexpression by endpoint PCR using MyTaq HS Red Mix (Meridian Bioscience cat. BIO-25048) with primers AluJb_qPCR4 forward/reverse and these PCR settings: initial denaturation (95°C, 1 minute) and 35 cycles of denaturation (95°C, 15 seconds), annealing (60°C, 15 seconds), and extension (72°C, 10 seconds). We note that the primers used here were designed to specifically amplify the *AluJb* copy on the overexpression plasmid by targeting the *AluJb* 3’end and the downstream plasmid backbone sequence.

The integrity of RNA samples was evaluated using an Agilent High Sensitivity RNA ScreenTape assay (Agilent Technologies). Of the 5 biological replicates per group in the paired transcriptome/proteome experiment, the 4 samples with the highest eRIN scores, ranging between 6.2 and 8.8, were selected for sequencing. In the follow-up experiment with standard serum-containing media, all samples had an eRIN score > 9. We submitted total RNA samples to Novogene (Sacramento, California) for unstranded mRNA library preparation and sequencing on the NovaSeq 6000 platform as paired-end 150 bp reads. The raw FASTQ reads have been deposited to SRA under BioProject PRJNA1008694.

### Protein digestion and desalting

Frozen cellular pellets and frozen conditioned media were shipped to the USC-Buck Institute Nathan Shock Center Cellular Senescence and Beyond Core (CSBC) for proteomic analysis. Conditioned media (30 mL per sample) from either empty vector (N=4) or *Alu* overexpression (N=4) IMR-90 cultures were concentrated using 3 kDa molecular cut-off filters (Millipore Sigma, Burlington, MA) and protein concentrations were determined using Bicinchoninic Acid (BCA) assay (Thermo Fisher Scientific, Waltham, MA). Cell pellets from either empty vector (N=5) or *Alu* overexpression (N=5) IMR-90 cultures were solubilized in 100 µL of 0.5% sodium dodecyl sulfate (SDS) in 100 mM Triethylamonium bicarbonate (TEAB) with 1X protease inhibitor cocktail (PIC). Protein concentrations were determined using BCA assay.

Samples originating from cell pellets (∼100 µg) were brought to the same overall volume of 100 µL with water, while secretome samples (∼100 µg) were prepared directly. All samples were reduced using 20 mM dithiothreitol in 50 mM TEAB at 50°C for 10 minutes, cooled to room temperature (RT) and held at RT for 10 minutes, and alkylated using 40 mM iodoacetamide in 50 mM TEAB at RT in the dark for 30 minutes. Samples were acidified with 12% phosphoric acid to obtain a final concentration of 1.2% phosphoric acid. S-Trap buffer consisting of 90% methanol in 100 mM TEAB at pH ∼7.1 was added and samples were loaded onto S-Trap mini spin columns (Protifi). The entire sample volume was spun through the S-Trap mini spin columns at 4,000 x g and RT, binding the proteins to the micro spin columns. Subsequently, S-Trap micro spin columns were washed twice with S-Trap buffer at 4,000 x g and RT and placed into clean elution tubes. Samples were incubated for one-hour at 47°C with sequencing grade trypsin (Promega, San Luis Obispo, CA) dissolved in 50 mM TEAB at a 1:25 (w/w) enzyme:protein ratio. An additional aliquot of trypsin dissolved in 50 mM TEAB was added and samples were digested overnight at 37°C.

Peptides were sequentially eluted from mini S-Trap spin columns with 50 mM TEAB, 0.5% formic acid (FA) in water, and 50% acetonitrile (ACN) in 0.5% FA. After centrifugal evaporation, samples were resuspended in 0.2% FA in water and desalted with Oasis 10-mg Sorbent Cartridges (Waters, Milford, MA). The desalted elutions were then subjected to an additional round of centrifugal evaporation and re-suspended in 0.1% FA in water at a final concentration of 1 µg/µL. Eight microliters of each sample were diluted with 2% ACN in 0.1% FA to obtain a concentration of 400 ng/µL. One microliter of indexed Retention Time Standard (iRT, Biognosys, Schlieren, Switzerland) was added to each sample, thus bringing up the total volume to 20 µL [32].

### Mass spectrometric analysis (secretome)

Reverse-phase high-performance liquid chromatography (HPLC)-MS/MS analyses were performed on a Dionex UltiMate 3000 system coupled online to an Orbitrap Exploris 480 mass spectrometer (Thermo Fisher Scientific, Bremen, Germany). The solvent system consisted of 2% ACN, 0.1% FA in water (solvent A) and 80% ACN, 0.1% FA in ACN (solvent B). Digested peptides (800 ng) were loaded onto an Acclaim PepMap 100 C_18_ trap column (0.1 x 20 mm, 5 µm particle size; Thermo Fisher Scientific) over 5 minutes at 5 µL/min with 100% solvent A. Peptides were eluted on an Acclaim PepMap 100 C_18_ analytical column (75 µm x 50 cm, 3 µm particle size; Thermo Fisher Scientific) at 300 nL/min using the following gradient: linear from 2.5% to 24.5% of solvent B in 125 min, linear from 24.5% to 39.2% of solvent B in 40 min, up to 98% of solvent B in 1 min, and back to 2.5% of solvent B in 1 min. The column was re-equilibrated for 30 min with 2.5% of solvent B, and the total gradient length was 210 min. Each sample was acquired in data-independent acquisition (DIA) mode [33–35]. Full MS spectra were collected at 120,000 resolution (Automatic Gain Control (AGC) target: 3e6 ions, maximum injection time: 60 ms, 350-1,650 *m/z*), and MS2 spectra at 30,000 resolution (AGC target: 3e6 ions, maximum injection time: Auto, Normalized Collision Energy (NCE): 30, fixed first mass 200 *m/z*). The isolation scheme consisted of 26 variable windows covering the 350-1,650 *m/z* range with an overlap of 1 *m/z* (see **Supplementary Table S2A**) [34].

### Mass spectrometric analysis (intracellular protein)

LC-MS/MS analyses were performed on a Dionex UltiMate 3000 system coupled to an Orbitrap Eclipse Tribrid mass spectrometer (both from Thermo Fisher Scientific, San Jose, CA). The solvent system consisted of 2% ACN, 0.1% FA in water (solvent A) and 98% ACN, 0.1% FA in water (solvent B). Proteolytic peptides (600 ng) were loaded onto an Acclaim PepMap 100 C_18_ trap column (0.1 x 20 mm, 5-µm particle size; Thermo Fisher Scientific) for 5 min at 5 µL/min with 100% solvent A. Peptides were eluted on an Acclaim PepMap 100 C_18_ analytical column (75 µm x 50 cm, 3 µm particle size; Thermo Fisher Scientific) at 300 nL/min using the following gradient of solvent B: 2% for 5 min, linear from 2% to 20% in 125 min, linear from 20% to 32% in 40 min, up to 80% in 1 min, 80% for 9 min, and down to 2% in 1 min. The column was equilibrated with 2% of solvent B for 29 min, with a total gradient length of 210 min. All samples were acquired in DIA mode. Full MS spectra were collected at 120,000 resolution (AGC target: 3e6 ions, maximum injection time: 60 ms, 350-1,650 m/z), and MS2 spectra at 30,000 resolution (AGC target: 3e6 ions, maximum injection time: Auto, NCE: 27, fixed first mass 200 m/z). The isolation scheme consisted of 26 variable windows covering the 350-1,650 m/z range with an overlap of 1 *m/z* (see **Supplementary Table S2A**) [34].

### RNA-seq read trimming, mapping, and quantification

Fastq files were first trimmed using fastp v0.20.1 [36] to (i) remove adapter sequences, (ii) hard trim the first 15 base pairs of each read to remove biased sequence composition, (iii) remove low quality bases, and (iv) remove reads shorter than 36 base pairs. Read quality for each sample was then inspected using fastqc v0.11.9. Next, the GRCh38 primary human genome assembly and comprehensive gene annotation were obtained from GENCODE release 44 [37]. We modified the reference genome to include the sequence for the *AluJb* overexpression plasmid as an additional contig. The trimmed reads were aligned to this modified reference genome using STAR v2.7.3a [38] with the following parameters: outFilterMultimapNmax 100, winAnchorMultimapNmax 100, and outFilterMismatchNoverLmax 0.04. Finally, the TEcount function in the TEtranscripts v2.1.4 [39] package was employed to obtain gene and repeat subfamily counts, using the GENCODE annotations to define gene boundaries and a repeat GTF file provided on the Hammell lab website (downloaded on September 14 2023 from https://labshare.cshl.edu/shares/mhammelllab/www-data/TEtranscripts/TE_GTF/GRCh38_GENCODE_rmsk_TE.gtf.gz) to define repeat element boundaries.

### Differential gene expression analyses

Gene and repeat subfamily count files were loaded into R v4.3.3. To filter lowly expressed genes in each analysis, a counts-per-million (cpm) threshold corresponding to 10 reads in the median-length library was defined. Genes and repeat subfamilies were kept for analysis if they were expressed at levels surpassing this cpm threshold in at least as many samples as the smallest group. DESeq2 v1.42.1 [40] was used to identify significant (FDR < 0.05) differentially expressed genes and repeat subfamilies between groups. To visualize sample grouping patterns from the expression data, we carried out multidimensional scaling (MDS) analysis using a distance metric between samples based on Spearman’s rank correlation value (1-Rho), which was then provided to the ‘cmdscale’ R function.

### Re-analysis of publicly available transcriptomic data

We leveraged publicly available mRNA-sequencing data for human primary fibroblasts derived from 133 healthy aging individuals [41] to characterize the effects of aging on the repetitive element transcriptome. Though others have conducted similar analyses [42, 43], we applied additional sample processing and filtering criteria to remove batch effects and isolate aging effects. We only kept samples from adults at least 20 years old to avoid the confounding effects of human development occurring prior to adulthood. We focused on samples derived from people who specified ‘Caucasian’ for their ethnicity to avoid confounding factors due to rare genotypes, since this was the most well-represented group. We only retained fibroblast samples extracted from skin on the arm to avoid potential confounding effects from extraction site. Thus, from an initial 133 samples, we obtained a filtered, curated set of 82 samples for downstream analysis. Fastq files for the curated set of samples were trimmed, mapped, and quantified as described above, with three modifications: (i) since raw reads were of two different lengths, the longer reads were hard trimmed with fastp v0.20.1 [36] at the 3’ end to ensure that read lengths were equal across all samples, (ii) reads were mapped to the unmodified human reference genome, and (iii) individual repeat locus counts, in addition to the repeat subfamily counts, were generated using the TElocal v1.1.1 package and a repeat file provided on the Hammell lab website (downloaded on October 31 2023 from https://labshare.cshl.edu/shares/mhammelllab/www-data/TElocal/prebuilt_indices/). Individual repeat locus counts were then aggregated by subfamily and the following genomic context categories as previously described [44]: intronic, exon-overlapping, intergenic and near a gene (within 5 kilobases), or intergenic and distal from a gene (farther than 5 kilobases).

With the generated counts files, we then checked for potential sample mislabeling or cross-contamination by checking the expression of the sex-specific markers *XIST* and *DDX3Y*. Six potential outliers using the 1.5*IQR (interquartile range) rule were omitted from downstream analyses. This yielded 76 samples that were utilized for differential gene expression analysis across age, which was carried out as listed above with these alterations: (i) a gene was considered expressed if it met the cpm threshold in at least 20% of samples and (ii) batch effects corresponding to biological sex and sequencing instrument were regressed out from the raw read counts using the ‘removeBatchEffect’ function in the R package Limma v3.58.1 [45].

### DIA-MS Data Processing and Statistical Analysis

DIA data was processed in Spectronaut (versions 14.10.201222.47784 and 15.1.210713.50606) using directDIA. Data extraction parameters were set as dynamic and non-linear iRT calibration with precision iRT was selected. Data was searched against the *Homo Sapiens* reference proteome with 20,380 entries (UniProtKB-SwissProt), accessed on 01/29/2021. Trypsin/P was set as the digestion enzyme and two missed cleavages were allowed. Cysteine carbamidomethylation was set as a fixed modification while methionine oxidation and protein N-terminus acetylation were set as dynamic modifications. Identification was performed using 1% precursor and protein q-value. Quantification was based on the peak areas of extracted ion chromatograms (XICs) of 3 – 6 MS2 fragment ions, specifically b- and y-ions, with q-value sparse data filtering and iRT profiling applied (**Supplementary Table S2F**). Local normalization was applied for intracellular protein analysis (i.e., cell pellets), but not for secretome analysis. Differential protein expression analysis comparing 1) *Alu* overexpression to empty vector was performed using a paired t-test, and p-values were corrected for multiple testing, using the Storey method [46]. Specifically, group wise testing corrections were applied to obtain q-values. For intracellular protein analysis, protein groups with at least two unique peptides, q-value < 0.05, and absolute log_2_(fold-change) > 0.2 were called as significantly-altered (**Supplementary Table S2E**). For secretome analysis, protein groups with at least two unique peptides, q-value < 0.05, and absolute log_2_(fold-change) > 0.58 were called as significantly-altered (**Supplementary Table S2F**).

### Functional enrichment and network analyses

We used the Gene Set Enrichment Analysis (GSEA) paradigm as implemented in the R package clusterProfiler v4.10.1 [47]. Multi-contrast pathway enrichment was also carried out across “-omics” layers using the mitch v1.14.0 R package [48]. Gene Ontology (GO) gene sets were obtained from the Molecular Signature Database release 2024.1.Hs [49, 50]. Reactome v92 pathway gene sets were obtained directly from the Reactome website [51]. We also obtained several aging- and senescence-associated gene sets, including core senescence-associated secretory phenotype (SASP) factors [52], SenMayo [53], CellAge build 3 [54], aging-related genes derived from the Genotype-Tissue Expression project [55], and differentially expressed genes/repeats identified in our analysis of the human aging fibroblast data [41]. Finally, we obtained previously published lists of genes upregulated and downregulated by stable overexpression of potentially transposition-competent *AluSq2* or *AluSx* transposons in IMR-90 fibroblasts [56]. For simplicity, the union of upregulated genes and the union of downregulated genes across both conditions were defined as *AluS* upregulated and downregulated genes, respectively.

For gene set enrichment analysis of transcriptomic changes, the DESeq2 v1.42.1 Wald-statistic was used to generate a ranked list of genes and repeats. For gene set enrichment analysis of proteomic and secretomic changes, the product of the log_2_(fold change) and - log_10_(Q-value) was used to generate a ranked list of proteins, which were then mapped to their corresponding gene symbol. All gene sets with an FDR < 0.05 were considered significant. For plots with a single analysis, the top 5 downregulated and top 5 upregulated gene sets were plotted, at most. For plots with multiple analyses, shared gene sets with consistent expression patterns across individual analyses were first identified. Then, the p-values for shared gene sets were combined using Fisher’s method, and this meta-analysis p-value was used to rank shared gene sets. Finally, the top 5 upregulated gene sets and the top 5 downregulated gene sets were plotted, at most. If there were less than 5 gene sets in either group, those were replaced with gene sets exhibiting the opposite regulation, in order to plot 10 shared gene sets whenever possible. For multi-contrast pathway enrichment analysis with mitch, gene sets with an adjusted MANOVA (Multivariate Analysis of Variance) p-value < 0.05 were considered significant. For mosaic plots comparing the abundances of gene set members in differentially versus non-differentially expressed genes/proteins, statistical significance was assessed using Fisher’s method, and comparisons with p < 0.05 were considered significantly enriched.

For the transcriptional data, predicted transcription factor activity was inferred using the decoupleR v2.8.0 package [57]. The ‘fgsea’ algorithm was run on the list of genes, and their accompanying Wald statistic, using the CollecTRI curated collection of transcription factors and their targets [58]. All regulons with a p < 0.05 were considered significantly differentially impacted, and the top 5 upregulated regulons and the top 5 downregulated regulons by magnitude were plotted. In addition, a gene network was constructed with NetworkAnalyst [59–61] on May 15, 2025. All differentially expressed genes (FDR < 0.05) were ordered by their Wald statistic and submitted to NetworkAnalyst as a gene list. A network was then constructed by taking the union of a generic PPI (first order network using the STRING interactome database, 900 confidence score cutoff, and requiring experimental evidence) and a signaling network (using the SIGNOR 2.0 database). The most populated subnetwork, subnetwork 1, was then visualized on the platform.

### Functional Assays

For functional assays, 2-4*10^6^ IMR-90 fibroblasts were transfected by electroporation as previously described and allowed to recover for ∼24 hours on 6-well plates. Six independent transfections per group served as biological replicates in each experiment.

#### Metabolic profiling using the Agilent Seahorse platform

To assess respiratory capacity, cells were detached, and 90,000 cells per well were seeded on a Seahorse XF Pro M Cell Culture Microplate (Agilent Part No. 103774-100) containing maintenance media. For each independent transfection, three technical replicate wells were prepared and cultured for 24 hours. Then, cells were washed and cultured in unbuffered Seahorse XF base media (Agilent cat. 102353-100) containing 5.5 mM of glucose, and the Seahorse XF Cell Mito Stress Test (part no. 103015-100) was carried out by the USC Leonard Davis School of Gerontology Seahorse Core. Afterwards, the media was removed, cells were lysed by freezing at −80°C overnight, 100 μL of water were added, the plate was placed in a 37°C incubator for 1 hour, and cells were frozen at −80°C for another hour. After thawing, a 200 μg/mL working solution of Hoechst 33342 in Tris-NaCl-EDTA (TNE) buffer was prepared, and 100 μL were added per well. Fluorescence was measured using an excitation wavelength of 346 nm and an emission wavelength of 460 nm, in order to normalize for cell number differences across wells. This experiment was repeated twice which yielded n = 12 independent transfection samples per group, the values were normalized to the median of the control group in each experiment, the values across experiments were combined, and statistical significance was reached if a Wilcoxon test yielded p < 0.05.

#### Cell cycle stage profiling using propidium iodide staining

Cell cycle status was determined by propidium iodide (PI) staining. After the recovery period, cells were detached, and ∼500,000 - 600,000 cells per well were seeded on 6-well plates containing maintenance media. After 24 hours, cells were detached, cells were spun down (300xG, 5 minutes), cells were washed once in DPBS, cells were fixed by drop-wise adding pre-chilled 70% ethanol, and fixed cells were stored at −20°C for a minimum of 12 hours. Then, cells were spun down (500xG, 5 minutes, 4°C), cells were washed once with ice-cold DPBS, cells were resuspended in labeling buffer, and cells were stained for 30 minutes in the dark. Staining buffer was prepared in DPBS with the following final concentrations: 50 μg/mL propidium iodide (Alfa Aesar cat. J66584), 100 μg/mL PureLink RNase A (Invitrogen cat. 12091021), and 0.05% Triton X-100 (MP cat. 194854). This experiment was repeated three times which yielded n = 18 independent transfection samples per group, the values were normalized to the median of the control group in each experiment, the values across experiments were combined, and statistical significance was reached if a Wilcoxon test yielded p < 0.05.

#### Protein aggregation assessment using proteostat staining

The relative amounts of protein aggregates were determined using the PROTEOSTAT Aggresome Detection Kit (Enzo cat. No. ENZ-51035-K100). After the recovery period, cells were detached, and ∼500,000 - 600,000 cells per well were seeded on 6-well plates containing maintenance media. After 24 hours, the assay was carried out, preparing staining solution by adding Proteostat dye and Hoechst 33342 at a dilution of 1/10,000 and 1/1,000, respectively. This experiment was repeated four times which yielded n = 24 independent transfection samples per group, the values were normalized to the median of the control group in each experiment, the values across experiments were combined, and statistical significance was reached if a Wilcoxon test yielded p < 0.05.

Flow cytometry for Proteostat and cell cycle assays was carried out on a MACSQuant Analyzer 10 (Miltenyi Biotec, 130-096-343). Flow cytometry data was analyzed in Flowlogic Solution 1.0.

### Other software versions

Analyses were conducted using R version 4.3.3 and code was re-run independently on R version 4.3.3 to check for reproducibility. Flow cytometry results were analyzed using Flowlogic Solution 1.0.

#### Data and code availability

RNA-seq data have been deposited at the Sequence Read Archive (SRA) under BioProject PRJNA1008694. Raw data and complete mass spectrometry data sets have been uploaded to the Mass Spectrometry Interactive Virtual Environment (MassIVE) repository, developed by the Center for Computational Mass Spectrometry at the University of California San Diego, and can be downloaded using the following link: https://massive.ucsd.edu/ProteoSAFe/dataset.jsp?task=42e058c0f1244315aa9d3e61add4cdd3 (MassIVE ID number: MSV000097822; ProteomeXchange ID: PXD063739). Original DNA gel images, raw Agilent Seahorse data, and raw flow cytometry data have been deposited at Figshare at [DOI: 10.6084/m9.figshare.28928147]. All scripts used to analyze the data in this manuscript are available on the Benayoun Lab GitHub at [https://github.com/BenayounLaboratory/AluJB_fibroblast_overexpression].

## RESULTS

### Aging upregulates repetitive elements, including *Alu* transposons, in primary human fibroblasts

To systematically characterize the effects of aging on the repetitive transcriptome, we leveraged publicly available mRNA-sequencing data for human aging primary fibroblasts [41] (**Figure 1A**). Though others have explored the relationship between aging and transposon expression in these samples [42, 43], we note that our sample selection criteria were more stringent. To minimize batch effects and isolate aging effects without confounding effects from development, the final curated sample set we utilized included 76 samples with the following characteristics: “healthy” male and female adults were aged between 20 and 96 years old, adults were of Caucasian ethnicity, and all fibroblasts were derived from skin on the arm. Restricting the dataset to a demographically and anatomically homogenous sample subset is expected to minimize sample heterogeneity and to limit any spurious changes unrelated to aging that might confound downstream transcriptomic analyses (see methods).

**Figure 1.**
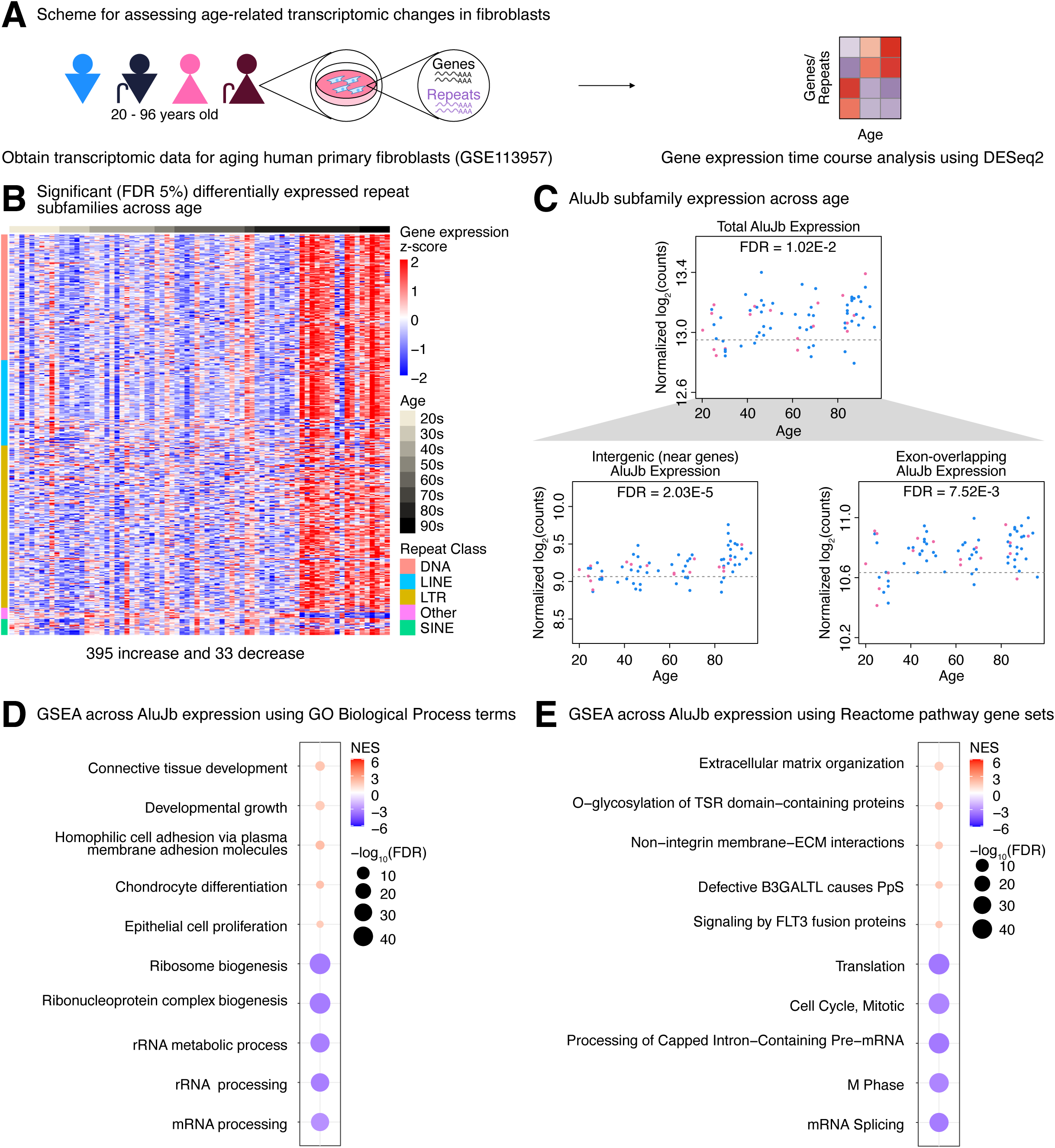
Aging upregulates repetitive elements, including *Alu* transposons, in primary human fibroblasts. **(A)** A diagram illustrating the transcriptomic time course analysis carried out using a publicly-available mRNA-sequencing dataset (GSE113957) for N = 76 aging primary human fibroblasts. **(B)** Gene expression heatmap for significant (FDR < 0.05) differentially expressed repeat subfamilies across age. **(C)** Scatterplots for the library size-normalized counts across age for the total *AluJb* subfamily expression, the *AluJb* subfamily expression from intergenic loci within 5 kilobases from a gene, and the *AluJb* subfamily expression from loci overlapping gene exons. All three differential expression results were significant at an FDR < 0.05. Females are represented by pink dots, and males are represented by blue dots. Differential gene expression analysis across total endogenous *AluJb* expression was carried out, followed by gene set enrichment analysis, and the top 5 significant (FDR < 0.05) **(D)** GO Biological Process and **(E)** Reactome pathway gene sets were plotted. FDR: False Discovery Rate.

Differential gene expression analysis identified 395 repeat subfamilies that increased with age and 33 subfamilies that decreased with age at an FDR < 0.05 (**Figure 1B, Supplementary Table S1A**). These effects were not specific to any repeat class, as DNA transposons, long interspersed elements (LINEs), short interspersed elements (SINEs), and long terminal repeat (LTR) transposons changed with age (**Supplementary Table S1A**). Because SINEs are a relatively understudied class of transposons despite being one of the most abundant classes in the human genome, we chose to focus on one of its subfamilies and characterize it further. Though a few SINE subfamilies changed with age, we decided to concentrate on the *AluJb* subfamily of retrotransposons, whose total expression increased with age (FDR = 1.02E-2, **Figure 1C**). Interestingly, younger *Alu* elements from the *AluS* and *AluY* subfamilies were more variable, although some members were significantly, but more modestly, upregulated (**Figure S1A-B; Supplementary Table S1A**). After quantifying expression levels at individual repeat loci and aggregating counts by genomic context [as previously done in [44]] (**Supplementary Table S1B**), we observed that intergenic *AluJb* copies within 5 kilobases of a gene (FDR = 2.03E-5) and exon-overlapping copies (FDR = 7.52E-3) both increased with age (**Figure 1C).** Intergenic *AluJb* expression increased at 40 years of age and continued increasing afterwards, whereas exonic *AluJb* expression rose around 40 years of age but then plateaued. Although *AluJb* copies showed distinct expression trajectories, observed patterns are consistent with the accepted notion that transposon control mechanisms become dysregulated with age including for *Alu* transposons, which can become elevated with age. Moreover, these results leave open the possibility that strongly upregulated Alu elements, such as *AluJb,* may exert cellular aging-associated effects either through its own independent transcripts or by incorporating itself into the transcripts of hosting genes.

To identify candidate pathways that may become mis-regulated in response to elevated *AluJb* expression, we carried out differential gene expression analysis across total *AluJb* expression levels (**Supplementary Table S1C**) followed by gene set enrichment analysis (GSEA). Interestingly, top Gene Ontology (GO) Biological Process terms associated with proteostasis, such as ‘ribosome biogenesis’, ‘rRNA metabolic process’, and ‘rRNA processing’, were significantly downregulated (**Figure 1D, Supplementary Table S1D**). Other terms associated with mitochondrial metabolism (like ‘aerobic respiration’ and ‘electron transport chain’) and the cell cycle (like ‘mitotic nuclear division’ and ‘cell cycle phase transition’) were also significantly downregulated. In contrast, top terms associated with the extracellular matrix, such as ‘homophilic cell adhesion via plasma membrane adhesion molecules’, were significantly upregulated. In parallel, top Reactome pathways associated with the cell cycle, such as ‘cell cycle, mitotic’ and ‘M phase’, as well as pathways associated with proteostasis, such as ‘translation’, were downregulated (**Figure 1E, Supplementary Table S1E**). Similar to the GO analysis, extracellular matrix-related terms such as ‘extracellular matrix organization’ were significantly upregulated. These results highlight recurrent correlations between *AluJb* expression and alterations in aging-associated pathways related to proteostasis, mitochondrial function, the cell cycle, and the extracellular environment, and raise the intriguing possibility that *AluJb* expression itself may drive these phenotypes.

### *AluJb* promotes widespread transcriptional and proteomic remodeling

Our analysis identified numerous pathways correlated with *AluJb* expression levels in aging fibroblasts, though it was unclear whether *AluJb* was an upstream driver of changes in these pathways or whether it was a simple bystander. To begin to answer this question and characterize the effects of elevated *AluJb* expression on primary cell phenotypes, we placed the consensus sequence under the control of the U6 RNA polymerase III promoter and transiently transfected human IMR-90 fibroblasts (**Figure 2A**). After a ∼24 hours recovery period following electroporation, cells were re-seeded at equal densities and cultured in low serum media in accordance with previously described guidelines for generating conditioned media from senescent cells and quiescent control cells [31]. We implemented this protocol because (1) we hypothesized that *Alu* overexpression may induce senescence-associated changes and (2) this protocol utilizes serum deprivation to induce quiescence in order to isolate senescent versus non-senescent effects and eliminate non-proliferating vs proliferating effects. After 72 hours, cells and conditioned media were collected for transcriptomic, proteomic, and secretomic analysis (**Figure 2A, Supplementary Tables S2A-C**). Importantly, overexpression of plasmid-specific *AluJb* was verified by reverse transcription (RT) followed by endpoint polymerase chain reaction (PCR) (**Figure S2A**). We chose to validate *AluJb* expression using end point PCR rather than quantitative PCR, as the empty vector control is expected to lack any amplification and thus cannot serve as a valid reference for RT-qPCR quantification. To note, we observed clear bands at the expected size in all our samples, although intensity varied between samples, consistent with successful *AluJb* overexpression (**Figure S2A**).

**Figure 2.**
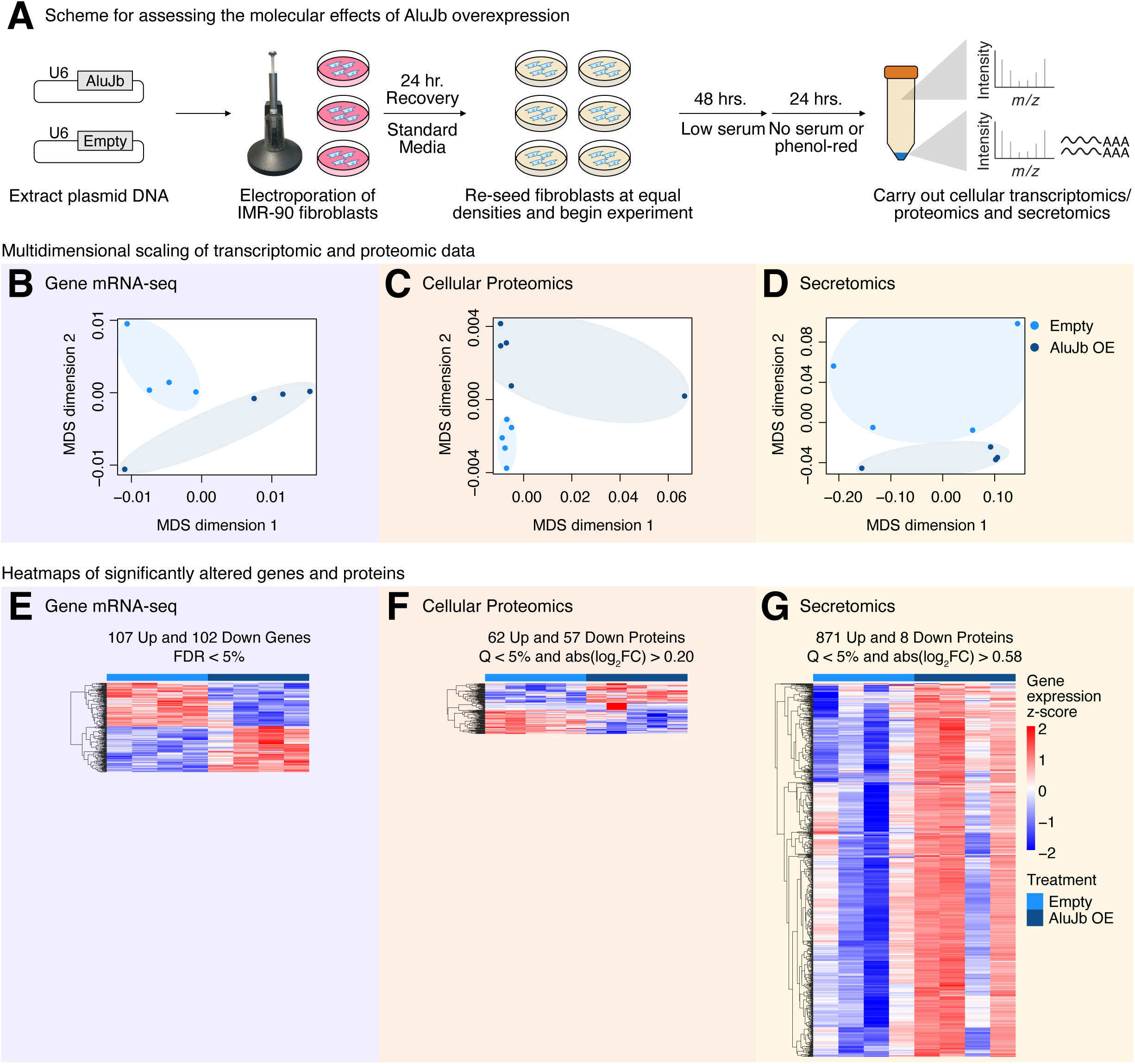
*AluJb* promotes widespread transcriptional and proteomic remodeling. **(A)** A diagram illustrating how control and *AluJb* overexpressing IMR-90 fibroblast samples were prepared for multi-omic profiling. N = 5 replicates were independently transfected per group, and N = 4 replicates were analyzed by mRNA-sequencing, N = 5 replicates were analyzed by cellular proteomic profiling, and N = 4 replicates were analyzed by secretome profiling. Multidimensional scaling (MDS) analysis of the **(B)** genic transcriptome, **(C)** cellular proteome, and **(D)** secretome. Abundance heatmaps for significant **(E)** differentially expressed genes (FDR < 0.05), **(F)** differentially abundant cellular proteins (Q < 5% and abs(log_2_FC) > 0.20), and **(G)** differentially secreted proteins (Q < 5% and abs(log_2_FC) > 0.58). FDR: False Discovery Rate, Q: Q-value. See also Figure S2 and S3.

Interestingly, multidimensional scaling (MDS) analysis segregated the empty vector control cells from the *AluJb*-overexpressing cells across the transcriptome (**Figure 2B**), cellular proteome (**Figure 2C**), and secretome (**Figure 2D**), though the profiles show in-group variation as expected from biological replicates. This suggests that *AluJb* overexpression induces major changes across “-omics” layers. Consistent with this notion, *AluJb* overexpression induced a significant (FDR < 0.05) upregulation of 107 genes and a downregulation of 102 genes (**Figure 2E, Supplementary Table S2D**), a significant (Q < 5% and abs(log_2_FC) > 0.20) upregulation of 62 cellular proteins and a downregulation of 57 cellular proteins (**Figure 2F, Supplementary Table S2E**), and a significant (Q < 5% and abs(log_2_FC) > 0.58) upregulation of 871 secreted proteins and a downregulation of 8 secreted proteins (**Figure 2G, Supplementary Table S2F**). Additionally, though the segregation was not as strong as with the gene expression profiles, the repeat subfamily expression profiles also segregated empty vector and *AluJb*-overexpressing cells by MDS analysis (**Figure S2B**), suggesting major shifts in the repeat transcriptome itself following *AluJb* overexpression. In line with this notion, we observed a trend where *AluJb*- overexpressing cells had more reads mapping to repeat subfamilies compared to control cells (**Figure S2C**). Moreover, the differential expression analysis identified 104 repeat subfamilies with significantly (FDR < 0.05) higher expression and 1 repeat subfamily with significantly lower expression in the overexpression group (**Figure S2D, Supplementary Table S2D**). These repeat subfamilies belonged to several repeat classes, including DNA transposons, LINEs, LTRs, and SINEs. Thus, these results suggest that elevated *AluJb* expression can promote the de-repression of other repeats from diverse repeat classes, potentially forming positive feedback loops where newly upregulated repeats further promote the expression of additional repeats.

Importantly, to contextualize our results within the broader *Alu* research literature, we compared our findings to a previous study of IMR-90 fibroblasts stably overexpressing two *AluS* retrotransposons [56] (**Figure S3A**). We observed that genes upregulated by *AluS* were significantly enriched among genes significantly upregulated by *AluJb* (Fisher’s exact p-value = 3.03E-3), and genes downregulated by *AluS* were significantly enriched among genes significantly downregulated by *AluJb* (Fisher’s exact p-value = 1.52E-6; **Figure S3B**). These results demonstrate a degree of consistency between *AluS-* and *AluJb*-regulated genes. As an alternative approach, we also generated gene sets for *AluS* induced and *AluS* repressed genes and carried out gene set enrichment analysis (GSEA) among *AluJb*-driven transcriptomic changes (**Figure S3C, Supplementary Table S2G**). Surprisingly, while the set of *AluS* induced genes was consistently upregulated by *AluJb*, the set of *AluS* repressed genes was inconsistently upregulated by *AluJb*. These results suggest that differences in the *Alu* subfamily (*AluJb* vs *AluS*) or differences in the overexpression duration (transient vs stable) may induce partially unique cellular responses and phenotypic effects.

### *AluJb* overexpression elicits aging-associated molecular alterations

Transposon de-repression is often seen in aging, but it is unclear whether aging follows transposon de-repression, or whether TE de-repression follows aging. To test the former idea, we compared our multi-omics measurements to published lists of senescence-induced or aging-induced gene changes (**Figure 3A**). Since our experimental approach was originally designed to capture senescence-associated secretory phenotype (SASP) factors [31], we first compared our multi-omics results to a list of core SASP factors [52]. Indeed, core SASP factors were enriched (p = 2.19E-2) among significant, rather than non-significant, differentially expressed genes (**Figure 3B**). Additionally, there was a suggestive enrichment of core SASP factors among significant, rather than non-significant, differentially abundant cellular or secreted proteins (**Figure 3C-3D**). These results suggest that *AluJb* may drive differences in core SASP factors across the transcriptome, cell proteome, and secretome without irreversible cell cycle arrest.

**Figure 3.**
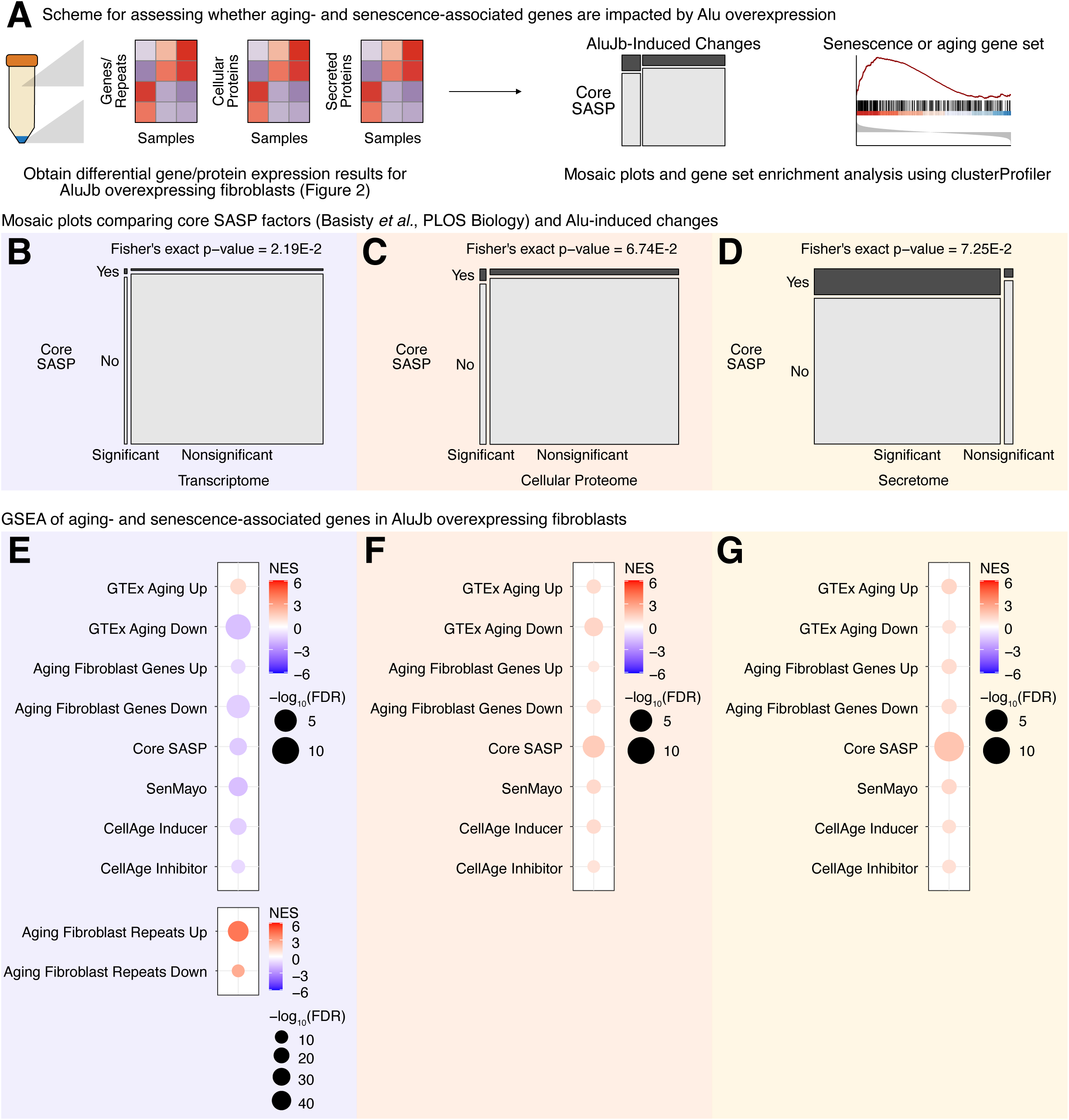
*AluJb* overexpression elicits aging-associated molecular alterations. **(A)** A diagram illustrating how *AluJb*-induced multi-omic changes and either senescence- or aging-induced features were compared. Mosaic plots showing the fraction of core SASP factors found among significant and non-significant *AluJb*-induced **(B)** mRNAs, **(C)** cellular proteins, and **(D)** secreted proteins. Statistical significance of frequency differences was assessed with Fisher’s exact test, and p < 0.05 was considered significant. GSEA analysis with gene sets for senescence- and aging-induced features in the *AluJb* **(E)** transcriptome, **(F)** cellular proteome, and **(G)** secretome. Gene sets with FDR < 0.05 were considered significant. SASP: Senescence-Associated Secretory Phenotype, FDR: False Discovery Rate, NES: Normalized Enrichment Score. See also Figure S4.

For heightened sensitivity and to determine the directionality of any observed changes, we carried out gene set enrichment analysis (GSEA) with several aging and senescence gene sets. At the transcriptomic level, we observed that (1) genes downregulated with age across tissues in the Genotype-Tissue Expression (GTEx) project [55], (2) genes downregulated with age in human primary fibroblasts [41], and (3) repetitive elements upregulated with age in human primary fibroblasts significantly (FDR < 0.05) changed with the same directionality following *AluJb* overexpression (**Figure 3E, Supplementary Table S3A**). Other gene sets like Senmayo [53] and the core SASP were significantly repressed following overexpression, while repeats that were downregulated in human aging primary fibroblasts were significantly upregulated following overexpression. At the cellular proteomics level, core SASP factors were significantly (FDR < 0.05) upregulated following *AluJb* overexpression (**Figure 3F, Supplementary Table S3B**). We observed that genes downregulated with age across tissues in the GTEx project were significantly upregulated at the protein level following overexpression, but the directionality was inconsistent. Inconsistent directionality may reflect misregulation of aging pathways in an acute fashion, which may have distinct directionality compared to long term, chronic, compensatory changes. Finally, at the secretomics level, core SASP factors were significantly (FDR < 0.05) upregulated following *AluJb* overexpression (**Figure 3G, Supplementary Table S3C**). These factors included insulin-like growth factor binding proteins (like IGFBP4 and IGFBP5), matrix metalloproteinases (like MMP1 and MMP2), and tissue inhibitors of metallopeptidase (like TIMP1). Though some of the effects vary in directionality, these results suggest that elevated *AluJb* expression can partially mirror molecular changes observed during aging. Additionally, the significant upregulation of core SASP factors in the secretome highlights *AluJb* as a potential driver of inflammation and altered intercellular communication.

We next tested the inverse hypothesis that transposon-mediated effects would follow chronological aging (**Figure S4A**). Surprisingly, *AluJb*-induced genes were not significantly enriched among significant primary fibroblast aging genes, and downregulated *AluJb* genes were actually significantly depleted compared to their frequency among nonsignificant aging downregulated genes (**Figure S4B**). GSEA highlighted an inverted regulation of *AluJb*-induced genes with aging, where *AluJb* downregulated genes were upregulated with age and *AluJb* upregulated genes were downregulated with age (**Figure S4C, Supplementary Table S3D**). In contrast, many repetitive elements that were upregulated following *AluJb* overexpression were also upregulated in aged primary human fibroblasts (**Figure S4D-S4E, Supplementary Table S3D**). Again, differences in directionality may stem from the transient nature of our transposon overexpression compared to the chronic upregulation of transposons with age. Nonetheless, these results demonstrate that aging partially recapitulates the effects of *AluJb* and the upregulation of many endogenous repetitive elements, in particular. Taken together with the observation that *AluJb* is sufficient to induce aging-associated changes, these results suggest the existence of a complex and non-linear feedback mechanism whereby (1) aging can promote heightened repetitive element expression, including upregulation of *<u>AluJb</u>*, (2) *AluJb* can promote alterations in aging pathways, including the upregulation of additional repetitive elements which may themselves alter aging pathways, creating feedback loop pressure, and (3) aging regulates chronic repetitive element-induced gene expression.

### *AluJb* drives molecular changes in mitochondrial, proteostasis, cell cycle, and extracellular matrix pathways

To assign functions to molecular changes caused by elevated *AluJb* expression, we carried out gene set enrichment analysis (GSEA) in each individual “-omics” layer using functional gene set collections (**Figure 4A**). In the transcriptome and using the Gene Ontology (GO) Biological Process gene set collection, we identified an upregulation of extracellular matrix-related processes (*e.g*. “homophilic cell adhesion via plasma membrane adhesion molecules” and “cell junction organization”), as well as cell cycle-related processes (*e.g*. ‘regulation of mitotic sister chromatid segregation’; **Figure 4B, Supplementary Table S4A**). We also observed a downregulation of terms related to mitochondrial metabolism (*e.g*. “aerobic respiration” and “generation of precursor metabolites and energy”) and proteostasis (*e.g*. “cytoplasmic translation”). In the cellular proteome, the top upregulated processes were related to the cell cycle (*e.g*. “cell cycle G2/M phase transition”) and to protein phosphorylation (*e.g*. “regulation of dephosphorylation”; **Figure 4C, Supplementary Table S4B**). Finally, in the secretome, the top upregulated terms were related to responding to stimuli (*e.g*. “response to endogenous stimuli” and “response to transforming growth factor beta”) and the top downregulated terms were related to aspects of proteostasis (*e.g*. “rRNA processing”; **Figure 4D, Supplementary Table S4C**). GSEA with Reactome pathway gene sets [51] mirror many of the results obtained using the GO Biological Process terms. In the transcriptome, we again observed an upregulation of extracellular matrix-related pathways (*e.g*. “extracellular matrix organization” and “integrin cell surface interactions”) and a downregulation of proteostasis pathways (*e.g*. “eukaryotic translation elongation” and “eukaryotic translation termination”) and mitochondrial metabolic pathways (*e.g*. “aerobic respiration and respiratory electron transport”; **Figure 4E, Supplementary Table S4D**). In this case, we could not detect significantly (FDR < 0.05) altered pathways in the cellular proteome (**Figure 4F**). However, the secretome also exhibited an upregulation of extracellular matrix-associated pathways, as well as a downregulation of pathways related to DNA repair or epigenetic modifications (*e.g*. “base-excision repair, AP site formation” and “chromatin modifications during the maternal to zygotic transition”; **Figure 4G, Supplementary Table S4E**). These results demonstrate that *AluJb* can regulate functional pathways that are often differentially regulated with age.

**Figure 4.**
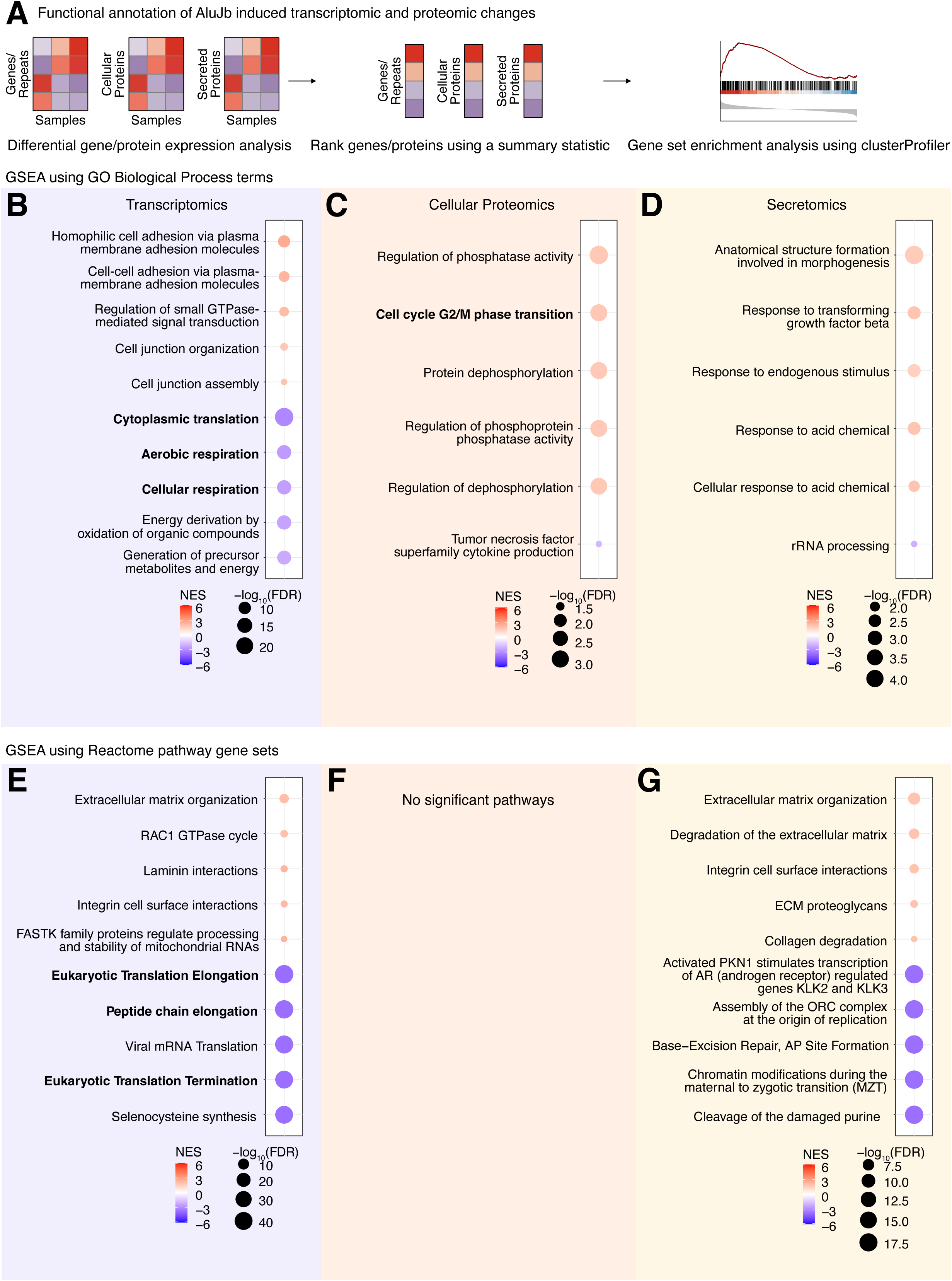
*AluJb* drives molecular changes in mitochondrial, proteostasis, cell cycle, and extracellular matrix pathways. **(A)** A diagram illustrating how functional annotations were assigned to transcriptomic and proteomic changes with gene set enrichment analysis. The top 5 significant (FDR < 0.05) GO Biological Process gene sets in the **(B)** transcriptome, **(C)** cellular proteome, and **(D)** secretome were plotted. The top 5 significant (FDR < 0.05) Reactome pathway gene sets in the **(E)** transcriptome, **(F)** cellular proteome, and **(G)** secretome were also plotted. FDR: False Discovery Rate, NES: Normalized Enrichment Score. See also Figure S5.

To ensure that these pathway-level changes were not unique to serum deprived cells and were also observed in proliferating cells (where more of these pathways could be functionally tested without the added stress of serum-deprivation), we carried out an additional transcriptomic analysis of proliferating IMR-90 fibroblasts cultured in standard rich media (**Figure S5A**). Again, we confirmed plasmid-specific *AluJb* overexpression by PCR (**Figure S5B**), observed a segregation of control and overexpression samples by MDS analysis of gene (**Figure S5C**) and repeat (**Figure S5D**) expression profiles, and observed a trending increasing in the percent of reads mapping to repeat subfamilies following *AluJb* overexpression (**Figure S5E**). We observed a significant (FDR < 0.05) increase in 3790 genes and a decrease in 987 genes, as well as an increase in 201 repeat subfamilies (**Figure S5F, Supplementary Table S4F**). The altered repeat subfamilies belonged to several repeat classes, including DNA transposons, LINEs, and LTRs. GSEA analysis with GO Biological Process gene sets highlighted an upregulation of cell cycle processes (*e.g*. “chromosome segregation” and “centriole assembly”) and a repression of mitochondrial metabolic processes (*e.g*. “aerobic respiration” and “oxidative phosphorylation”) and proteostasis processes (*e.g*. “protein folding”) (**Figure S5G, Supplementary Table S4G**). Likewise, GSEA with Reactome pathway gene sets highlighted a repression of proteostasis pathways (*e.g*. “eukaryotic translation elongation”, “peptide chain elongation”, and “HSF1 activation”) and mitochondrial metabolic pathways (**Figure S5H, Supplementary Table S4H**). These results demonstrate that *AluJb* can also promote broad molecular alterations in rich media conducive to cell proliferation.

Using the initial transcriptomic data, we next sought to characterize *AluJb*-induced alterations across broad gene modules (**Figure 5A**). An enrichment analysis of transcription factor (TF) regulons identified heat shock transcription factor 1 (HSF1) as one of the top TFs with repressed targets (**Figure 5B, Supplementary Table S4I**). Since HSF1 plays a crucial role in the heat shock response and therefore in maintaining proteostasis during stress [62], these results highlight potential master regulators of the previously noted *AluJb*-driven pathway alterations. Indeed, PRRX1 has been identified as a master transcription factor in stromal fibroblasts with functions related to extracellular remodeling [63], ETV5 regulates fatty acid metabolism through peroxisome proliferator-activated receptor (PPAR) signaling [64], and both KDM5A and CXXC1 play roles in epigenetic control [65, 66]. Additionally, a network analysis to construct a predicted protein-protein interaction (PPI) network using significantly differentially expressed genes highlighted two major clusters of corresponding proteins: one with proteins involved in mitochondrial function, such as *MT*-*ND2, MT-ND4, MT-ND5,* and *MT-ND6*, and one with proteins involved in translation, such as *RPS25* and *RPL36AL* (**Figure 5C**). These results further highlight broad-scale *AluJb*-driven alterations in molecular pathways involved with the extracellular environment, epigenetics, metabolism, and proteostasis.

**Figure 5.**
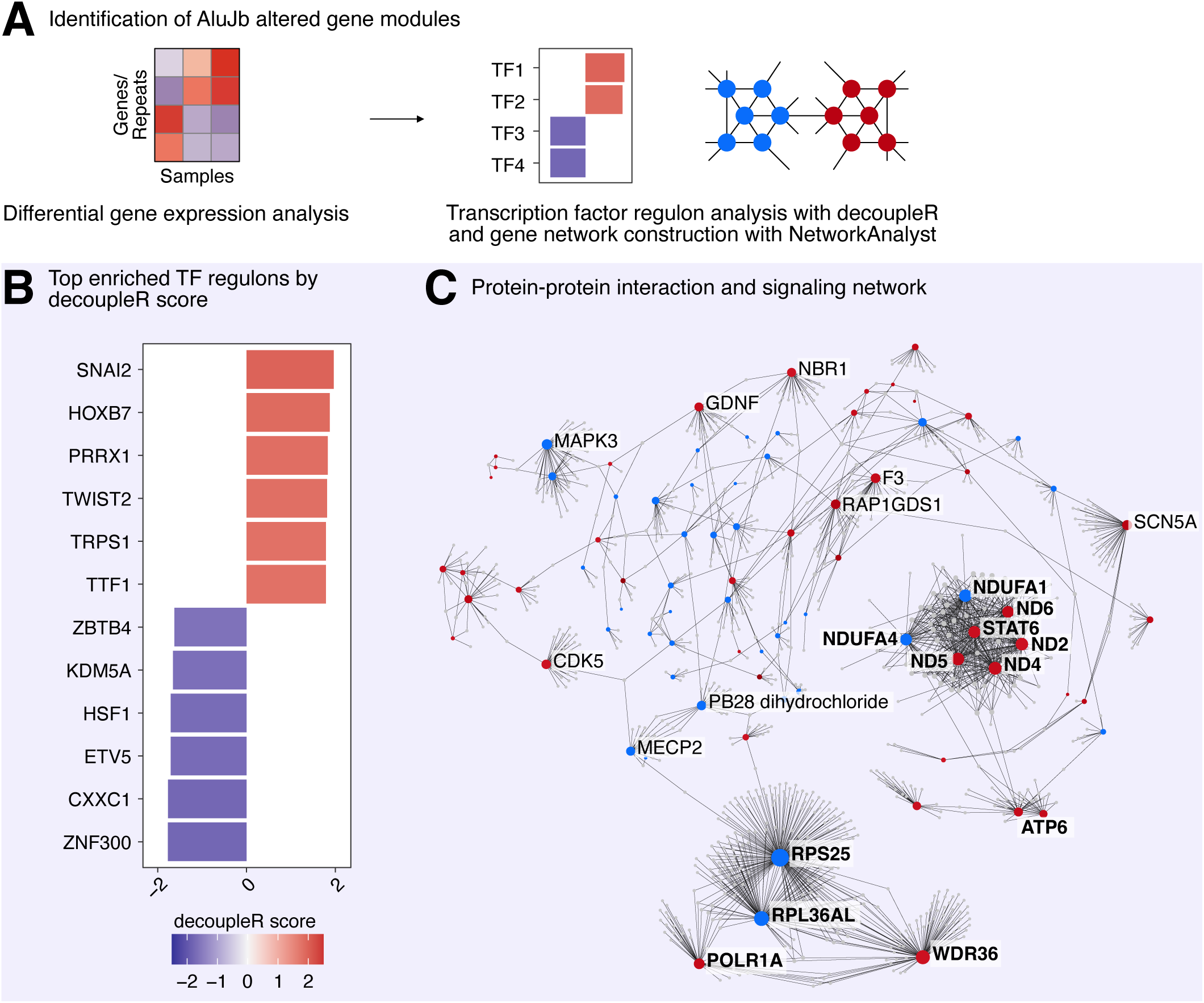
*AluJb* overexpression leads to disruption of transcription factor regulons and predicted PPI networks. **(A)** A diagram illustrating how the effects of *AluJb* on IMR-90 transcriptional networks was assessed. **(B)** Significantly (p < 0.05) enriched or depleted transcription factor regulons were identified with decoupleR, and the top 5 by magnitude were plotted. **(C)** Significant (FDR < 0.05) differentially expressed genes were used to construct a combined protein-protein interaction (PPI) and signaling network with NetworkAnalyst. The most populated subnetwork is shown. Blue nodes represent downregulated genes, and red nodes represent upregulated genes. FDR: False Discovery Rate.

### Multi-omic integration highlights alterations in extracellular, proteostasis, and metabolic pathways

To further increase sensitivity and identify pathways alterations occurring in more than one “-omics” analysis, we carried out multi-contrast gene set enrichment analysis (**Figure 6A**). Indeed, an overlap analysis of individual significant genes/proteins across “-omics” analyses was limited, and we identified four genes and corresponding proteins—*DCN, NES, POSTN*, and *DBI*—that changed in all three “-omics” layers (**Figure 6B**, **Figure S6A-S6C**). This modest overlap could result from multiple technical and biological reasons, including different detection sensitivities across “-omics” layers and imperfect correlations between “-omics” layers. In these cases, combinatorial analyses taking into account multiple “-omics” layers *a priori* (in the process of enrichment analysis) instead of *a posteriori* (overlapping differential results in each independent dataset) are more powerful to detect consistent functional changes elicited across layers.

**Figure 6.**
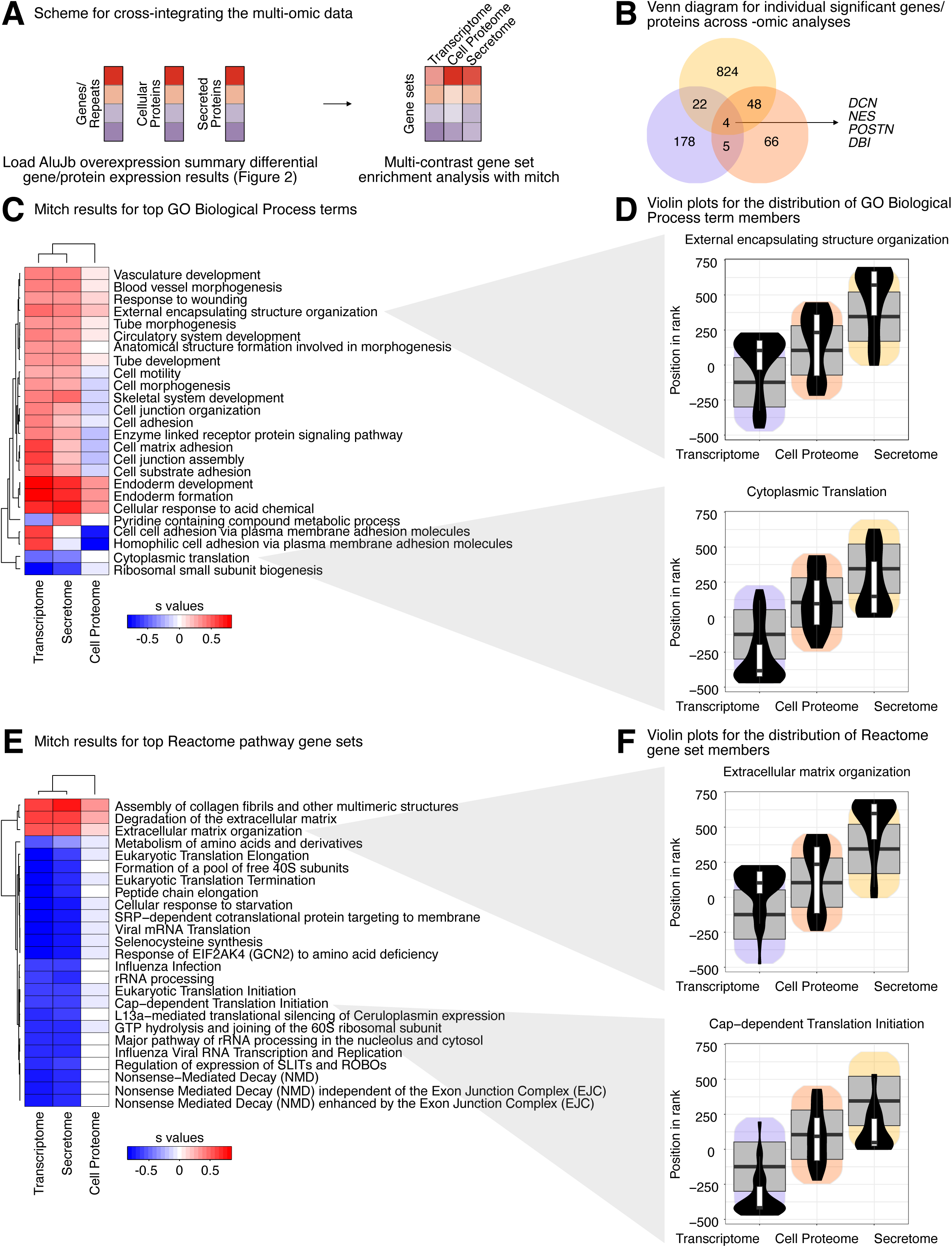
Multi-omics integration highlights alterations in extracellular, proteostasis, and metabolic pathways. **(A)** A diagram illustrating how transcriptome, proteome, and secretome changes were integrated via multi-contrast gene set enrichment analysis. **(B)** Venn diagram comparing individual genes and proteins that significantly changed in each “-omics” analysis. Highlighted are four genes and corresponding protein products—*DCN, NES, POSTN*, and *DBI*—that were altered in all three “-omics” layers. The top 25 multi-contrast gene set enrichment results using **(C)** GO Biological Process and **(E)** Reactome pathway gene sets. Gene sets with an adjusted MANOVA p-value < 0.05 were considered significant. Violin plots for select **(D)** GO Biological Process and **(F)** Reactome pathway gene sets related to recurring pathway themes are shown. See also Figure S6.

Thus, we performed multi-contrast gene set enrichment analysis using a rank MANOVA framework to identify pathways consistently regulated across our multi-omics profiling (**Figure 6**). We first leveraged GO Biological Process terms, which revealed that the transcriptome was most similar to the secretome, and that the cell proteome varied compared to the other two “-omics” types (**Figure 6C, Supplementary Table S5A**). This analysis also identified extracellular matrix-related terms, such as ‘external encapsulating structure organization’, ‘cell adhesion’, and ‘cell motility’, that were upregulated across the transcriptome and secretome but varied in the cell proteome (**Figure 6C-6D**). Similarly, proteostasis terms like ‘cytoplasmic translation’ and ‘ribosomal small subunit biogenesis’ were suppressed in the transcriptome and secretome but varied in the cell proteome (**Figure 6C-6D**). The multi-contrast analysis with the Reactome pathway gene sets also highlighted a stronger similarity between the transcriptome and secretome compared to the cell proteome (**Figure 6E, Supplementary Table S5B**). Mirroring the previous analysis, extracellular matrix terms like ‘extracellular matrix organization’ and ‘degradation of the extracellular matrix’ were upregulated across the three “-omics” analyses, while proteostasis terms like ‘cap-dependent translation initiation’ and ‘SRP-dependent cotranslational protein targeting to membrane’ were downregulated across all three “-omics” layers (**Figure 6E-6F**). These results further emphasize widespread alterations that are consistently observed in more than one “-omics” layer following *AluJb* overexpression.

### *AluJb* overexpression leads to functional alterations, as predicted from multi-omics alterations

Our individual and integrated multi-omics analyses identified molecular changes associated with several aging-regulated functional pathways, including mitochondrial metabolism, the cell cycle, and proteostasis, among others. However, whether these molecular changes translated to functional cellular changes remained unclear. To test the functional effects of *AluJb* on cell physiology, we thus carried out several functional assays (**Figure 7A**).

**Figure 7.**
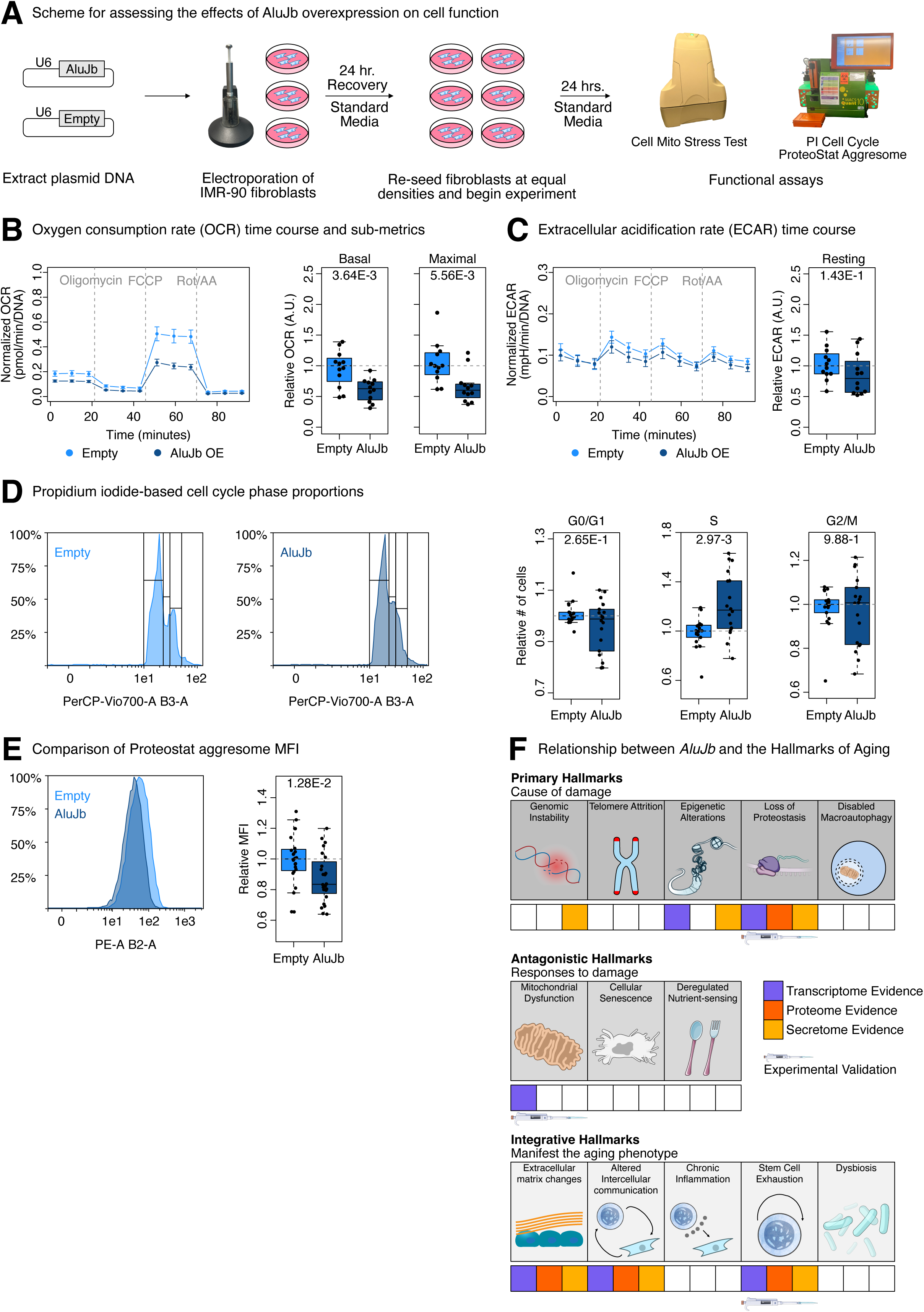
*AluJb*-induced functional alterations mirror molecular alterations and are consistent with disruption of aging hallmarks. **(A)** A diagram illustrating how the functional effects of *AluJb* overexpression were assessed. **(B)** A representative oxygen consumption rate time course from one experiment with N = 6 per group. Points represent means at each timepoint, and error bars represent the standard error of the mean. Two experiments, with a total of N = 12 per group, were used to calculate several OCR sub-metrics, including basal OCR and maximal OCR. Statistical significance was assessed with a Wilcoxon rank sum test, and p < 0.05 was considered significant. **(C)** A representative extracellular acidification rate time course from one of the aforementioned experiments. Points represent means at each timepoint, and error bars represent the standard error of the mean. Both experiments were used to calculate the resting ECAR, and statistical significance was assessed with a Wilcoxon rank sum test. **(D)** Representative flow cytometry histograms for propidium iodide-stained control and *AluJb* overexpressing cells. Three experiments were combined, for a total of N = 18 per group, and the relative number of cells in G0/G1, S, and G2/M phases in the control and overexpression group were compared with a Wilcoxon rank sum test. p < 0.05 was considered significant. **(E)** Representative flow cytometry histograms for Proteostat-stained control and *AluJb* overexpressing cells. Four experiments were combined, for a total of N = 24 per group, and the relative median fluorescence intensities between the control and overexpression groups were compared with a Wilcoxon rank sum test. p < 0.05 was considered significant. **(F)** A diagram summarizing the findings from this study as they relate to the hallmarks of aging. OCR: Oxygen Consumption Rate, ECAR: Extracellular Acidification Rate, MFI: Median Fluorescence Intensity. See also Figure S7.

First, we assessed mitochondrial respiration using a Seahorse Cell Mito Stress Test (**Figure 7B**). This revealed a significant (p < 0.05) reduction in many oxygen consumption rate (OCR)-related metrics, including basal respiration, maximal respiration, non-mitochondrial respiration, ATP-linked respiration, proton leak, and spare capacity (**Figure 7B**, **Figure S7A-S7B**). Though this assay is not specifically designed to evaluate resting glycolytic rate, parallel measurements of extracellular acidification rate did not reveal any significant differences in resting glycolytic rate (**Figure 7C**). These results demonstrate that mitochondrial respiration (but likely not glycolytic potential) is substantially dampened following *AluJb* overexpression, consistent with aging.

Next, we carried out a propidium iodide-based cell cycle assay (**Figure 7D, Figure S7C**). Since cell proliferation is one of a limited number of phenotypes known to be disrupted by *Alu* expression [56], we chose to re-analyze this phenotype since it (1) appeared in our analyses and (2) could serve as a potential positive control. Though we did not detect significant differences between control and *AluJb*-overexpressing cells in the G0/G1 and G2/M phases, we did detect a significant (p < 0.05) accumulation of *AluJb*-overexpressing cells in the S phase, consistent with a prior report [56]. However, contrary to this report, we interpret this accumulation in S-phase to represent likely replication stress rather than increased proliferative potential (based on total cell numbers over the course of our experiment, as well as the lack of a concomitant increase in G2/M cells). These results demonstrate a buildup of cells in the S phase, which may be caused by replication stress or stalling.

Finally, since we detected alterations in proteostasis-related pathways, we carried out a Proteostat Aggresome detection assay to identify differences in protein aggregate buildup (**Figure 7E, Figure S7D**). Interestingly, we observed a significantly (p < 0.05) lower median fluorescence intensity in the *AluJb*-overexpressing cells compared to control cells. These results demonstrate a reduced buildup of protein aggregates and further highlight alterations in proteostasis machinery.

Taken together, our molecular multi-omics analyses combined with functional profiling reveal *AluJb* retrotransposon-induced changes in several of the hallmarks of aging (**Figure 7F**) [67]. These include changes associated to (1) primary hallmarks like loss of proteostasis, epigenetic alterations, and genomic instability, (2) antagonistic hallmarks like mitochondrial dysfunction, and (3) integrative hallmarks like extracellular matrix changes, altered intercellular communication, and cell cycle changes that can manifest as stem cell exhaustion. Our results suggest that elevated *AluJb* retrotransposon expression is sufficient to drive mis-regulation of aging-related pathways and may thus directly contribute to aging phenotypes.

## DISCUSSION

### A resource for studying the effects of increased transposon expression in primary cells

In this study, we generated a multi-faceted resource for studying the effects of elevated *AluJb* transposon expression in “healthy” human IMR-90 fibroblasts. This resource encompasses a multi-omics characterization of the transcriptome, cell proteome, and secretome for human fibroblasts with and without *AluJb* overexpression. This resource expands the limited number of “-omics” resources where TE expression was directly modulated, including the previously mentioned transcriptomic study with stable *AluS* overexpression in IMR-90 fibroblasts [56], and a transcriptomic study with stably-integrated but inducible codon-optimized *LINE-1* expression in immortalized retinal pigment epithelial (RPE) cells [68]. Because RNA and protein levels do not always correlate [69], our study addresses molecular blind-spots present in prior studies focusing on a single “-omics” layer. Importantly, because aging involves the deterioration of healthy organs, tissues, and cells, it will be important (1) to identify and characterize the regulatory mechanisms that fail with age in healthy organs, tissues, and cells, including those that relate to transposon control, and (2) to characterize the effects of elevated transposon expression in healthy organs, tissues, and cells. In contrast to other studies utilizing cancer or immortalized cell lines to study transposons, our resource is primed for studying the causes and consequences of elevated transposon expression in “healthy”, non-cancerous, and non-immortalized cells. By integrating this resource with existing and newly-generated “-omics” resources where transposon expression is experimentally manipulated, it will be possible to characterize (1) organ-, tissue-, and cell type-specific changes in response to transposon expression, as well as (2) common and unique responses to different transposon families, subfamilies, or loci. Ultimately, we anticipate that this resource will be a useful starting point for answering these questions, especially as they relate to aging.

Our study also provides molecular signatures for elevated *AluJb* expression that can be readily incorporated into ongoing senescence, aging, therapeutic, or other studies with an interest in transposon control. Like the aging and senescence gene sets used to determine whether aging or senescence may be regulated by *AluJb*, the *AluJb* gene sets and protein sets we present can be used to determine whether transposon activation may be regulated by aging, senescence, or any other treatment of interest. As elevated transposon activity is a feature of many aging-associated diseases like cancer [12–15] and Alzheimer’s disease [16, 17], our molecular signatures can be used to simultaneously determine the effects of the disease and the effects of candidate therapeutics on transposon activity. Thus, we believe this resource holds promising potential for more thoroughly characterizing human diseases and the therapeutics used to treat them.

### Transposons as potential regulators of aging-related biological pathways

This study presents several observations that are consistent with the hypothesis that *AluJb* retrotransposons partially drive aging. First, we find that *AluJb* retrotransposons are upregulated in human aging primary fibroblasts, and their expression is correlated with alterations in pathways related to the hallmarks of aging [67], including pathways involved in mitochondrial metabolism, proteostasis, the cell cycle, and the extracellular matrix. Second, we find that *AluJb*-induced transcriptomic changes partially mirror transcriptomic changes in human aging primary fibroblasts and in aging tissues from the GTEx project. Moreover, both the cell proteome and secretome exhibited a significant upregulation of core SASP factors. Third, we find that functional pathways responding to *AluJb* overexpression are also related to the hallmarks of aging and include the pathways mentioned in the first point. Finally, we find that *AluJb* overexpression promotes differences in cell functions related to the hallmarks of aging. Our results add to the growing litany of studies demonstrating that *Alu* retrotransposons regulate features of aging.

Though the effects of *Alu* on cell function have not been broadly, unbiasedly, and extensively studied with “-omics” approaches, several targeted studies have explored *Alu*’s relationship to specific aging-regulated pathways, such as mitochondrial metabolism and the cell cycle. *Alu*’s influence on mitochondrial function has been explored in the context of advanced age-related macular degeneration. Specifically, in human cell culture and mouse models, *Alu* RNA was sufficient to promote opening of the mitochondrial permeability transition pore, release of mtDNA into the cytosol, and activation of inflammatory cGAS-STING signaling [70]. Whether this mechanism is specific to advanced age-related macular degeneration or is more broadly applicable to other aging contexts is an open question. Nonetheless, the mitochondrial dysfunction observed in that study is consistent with the repression of mitochondrial metabolism observed in this study.

With respect to cell cycle alterations, at least three studies have explored their relationship to *Alu* RNA levels. First, *Alu* transcripts were found necessary for senescence, as knockdown of *Alu* transcripts promoted exit of adult adipose-derived mesenchymal stem cells from senescence [10]. Second, consistent with this senescence-promoting model, introducing *Alu* RNA into RPE cells from humans with advanced age-related macular degeneration led to the upregulation of pro-inflammatory and senescence markers, including *IL-18, IL-1β, p16^INK4a^,* and SPiDER-*β*Gal staining [71]. Third, stable overexpression of two *AluS* elements in IMR-90 human fibroblasts was shown to upregulate cell cycle genes and induce an accumulation of cells in the S phase [56]. Though the authors speculated that the S phase accumulation reflects heightened cell proliferation, an alternative, more likely hypothesis is that it represents replication stress or stalling. Indeed, similar to this study, we also observe an accumulation of cells in the S phase without a concurrent accumulation in the G2/M phase, supporting the notion of replication stalling. Moreover, though we don’t detect a significant enrichment of senescence gene sets in any “-omics” layers, we do observe a significant upregulation of core SASP factors. Taken together, our study, combined with previously published studies, highlight the possibility that transcriptional upregulation of *Alu* transposons might be sufficient to regulate features of aging.

### Limitations of the study

While we believe that physiologically-relevant principles about *Alu* biology can be readily extracted from this study, we note potential further considerations. First, there are several viable options to choose from concerning any specific transposon to modulate, even when restricting the choice to a specific family, like *Alu*. We chose to focus on an *AluJb* element, instead of an *AluS* or *AluY* element, because (1) we observed that *AluJb* increased with age in human primary fibroblasts, making it a suitable candidate for studying its potential age-related effects and (2) we hypothesized that mobilization of an *AluJ* element would be much more restricted compared to an *AluS* or *AluY* element, potentially allowing us to focus on the effects of its transcriptional upregulation with minimal or reduced effects from mobilization. Additionally, we chose to use a consensus *AluJb* sequence instead of the sequence of a specific *AluJb* copy. We believe that the former approach can provide more generalizable results across *Alu* copies, and the latter approach would be better suited for identifying transposon locus-specific effects. Since we were interested in broad and conserved effects, we opted for the consensus sequence option. Additionally, we note that *AluJb* transfected cells produced comparable total RNA yield compared to empty vector control cells, indicating that observed effects are unlikely to stem from global changes in RNA biogenesis leading to acute transcriptional stress, though we cannot completely exclude that any similar-sized RNA of the same size might result in similar cellular changes.

Secondly, there are several options concerning the gene expression manipulation paradigm. We opted for a transient overexpression of *AluJb*, though stable *AluS* overexpression lines have been previously generated [56]. A comparison of the results from these two approaches suggests that they at least partially mirror each other (**Figure S3**). However, depending on whether one is interested in characterizing the effects of transient versus sustained *AluJb* expression, one approach may be better suited for the research question at hand. Although a stable overexpression approach may better model the sustained transposon expression observed during aging, “healthy” cells may have time to respond to the overexpression and employ compensatory regulatory mechanisms, such as DNA methylation, to restrict its impact. In addition, genetic drift occurring during selection may add noise to the analysis. This may or may not change the utility of the approach in modeling transposon-aging interactions. In the future, overexpression approaches could be complemented by the incorporation of knockdown approaches which can target endogenous *Alu* expression to provide additional insights. It is also worth mentioning that neither our short-term model, nor any single transposon activation event, can fully recapitulate the multifactorial biology of aging. The relationship between Alu activity and aging is likely complex and context-dependent alongside multiple interactions with various global/specific age-associated mechanisms such as DNA damage and/or chromatin remodeling. Hence, the divergences from established aging and senescence signatures may represent genuine, Alu specific responses that are independent from aging biology.

Finally, there are several options concerning the demographics underlying the samples being considered for experimentation. In this proof-of-principle study, we utilized a well-characterized female primary embryonic fibroblast cell line, IMR-90. As aging phenotypes, including TE expression, are frequently sex-dimorphic [72], it would be important to characterize both the common and sex-specific effects of *AluJb* overexpression, especially as they relate to aging. Additionally, though we used IMR-90 embryonic fibroblasts because of their frequent use as a model for studying senescence and aging, it would also be informative to assess the effects of *Alu* overexpression in adult primary fibroblasts, which may provide a more physiological environment to model the effects of elevated *Alu* expression with age. Ultimately, we believe this study serves as a useful starting point – a proof-of-principle - for the characterization of the relationship between transposon expression and aging.

## ACKNOWLEDGMENTS

This manuscript is based upon work supported by the National Science Foundation Graduate Research Fellowship Program (NSF GRFP) under Grant No. DGE-1842487 (to J.I.B.), the National Institute on Aging (NIA) under Grant No. T32 AG052374 (to J.I.B.), the University of Southern California with a Provost Fellowship (to J.I.B.), and the National Institute of General Medical Sciences (NIGMS) under Grant No. R35 GM142395 (to B.A.B). This work was also made possible by a pilot award under NIA grant P30 AG068345 (USC-Buck Institute Nathan Shock Center of Excellence) and by the NIH-OD instrumentation grant S10 OD028654 (to B.S.) for the Orbitrap Eclipse Tribrid mass spectrometry system.

We would like to thank Hemal Mehta at the USC Leonard Davis School of Gerontology Seahorse Core for conducting the seahorse assays. We are saddened by the passing of Dr. Judith Campisi and gratefully acknowledge her valuable contribution to this work before her passing.

Some artwork was taken from NIAID NIH BIOART Source (bioart.niaid.nih.gov) or from Bioicons (bioicons.com).

## AUTHOR CONTRIBUTIONS

**Juan I. Bravo:** Conceptualization, Data curation, Investigation, Formal analysis, Visualization, Writing – original draft (co-lead), Writing – review & editing. **Eyael Tewelde:** Investigation, Formal analysis, Writing – review & editing (co-lead). **Christina D. King:** Investigation, Formal analysis, Writing – original draft, Writing – review & editing. **Joanna Bons:** Investigation, Formal analysis, Writing – original draft, Writing – review & editing. **Samah Shah:** Investigation, Writing – review & editing. **Jacob Rose:** Investigation, Writing – review & editing. **Judith Campisi:** Conceptualization. **Birgit Schilling:** Conceptualization, Formal analysis, Writing – original draft, Writing – review & editing, Supervision, Funding acquisition. **Bérénice A. Benayoun:** Conceptualization, Data curation, Formal analysis, Visualization, Writing – original draft (co-lead), Writing – review & editing (co-lead), Supervision, Funding acquisition.

## DECLARATION OF INTERESTS

The authors declare that they have no competing interests. The content is solely the responsibility of the authors and does not necessarily represent the official views of the National Institutes of Health.

## Notes

### Competing Interest Statement

The authors have declared no competing interest.

### Summary of Updates

Clarifications of methodology and sample selection from the public fibroblast RNA-seq dataset; clarification of passage number for cells used; grammatical and syntactic edits.

