## Supplementary figures and legends for "A multi-omics analysis of human fibroblasts overexpressing an *Alu* transposon reveals widespread disruptions in aging-associated pathways"

**SUPPLEMENTARY MATERIALS**

**SUPPLEMENTARY FIGURES AND FIGURE LEGENDS**

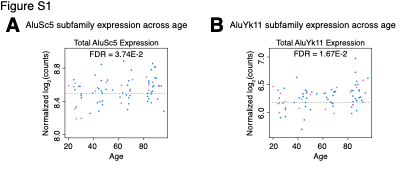

**Supplementary Figure S1. Aging upregulates diverse Alu subfamilies in primary human fibroblasts.** Scatterplots for the library size-normalized counts across age for **(A)** *AluSc5* and **(B)** *AluYk11* expression. Pink = Female, Blue = Male.

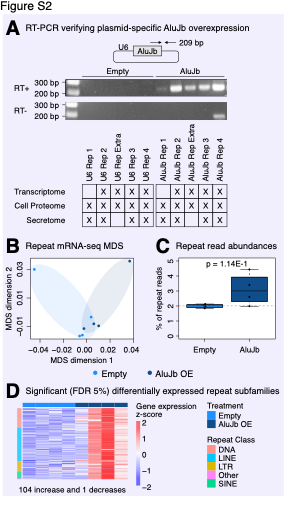

**Supplementary Figure S2. Endogenous repetitive elements are upregulated following *AluJb* overexpression.** **(A)** Plasmid-specific *AluJb* overexpression was assessed by endpoint RT-PCR of empty vector and *AluJb* overexpressing IMR-90 fibroblasts using primers targeting the 3’ *AluJb*-plasmid backbone junction. PCR reactions were carried out with (RT+) or without (RT-) reverse transcription in N = 5 samples per group, and N = 4-5 of these were used for multi-omic profiling. We note that the band on the lower right corner of the gel appears to be smaller in size than the expected amplicon size and may correspond to primer dimers. A table showing samples utilized for each -omics analysis is also shown. **(B)** Multidimensional scaling (MDS) analysis of the repetitive element transcriptome across samples. **(C)** Quantification of the percent of total reads mapping to repetitive elements. Statistical significance was assessed with a Wilcoxon rank sum test. **(D)** A gene expression heatmap for significant (FDR < 0.05) differentially expressed repeat subfamilies. RT: Reverse Transcription, FDR: False Discovery Rate.

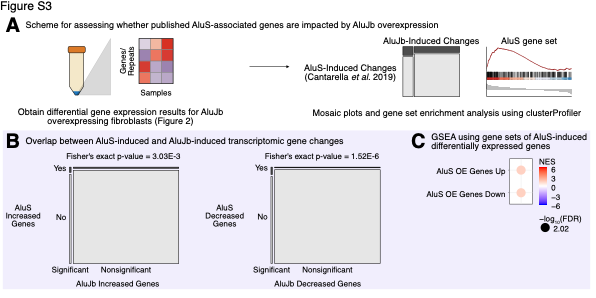

**Supplementary Figure S3. *AluJb*-induced transcriptomic changes partially mirror *AluS*-induced changes. (A)** A diagram illustrating how *AluJb*-induced and previously published *AluS*-induced transcriptomic changes were compared. **(B)** Mosaic plots showing the fraction of *AluS*-induced genes found among significant (FDR < 0.05) and non-significant *AluJb*-induced genes. Statistical significance of frequency differences was assessed with Fisher’s exact test, and p < 0.05 was considered significant. **(C)** GSEA analysis with gene sets for *AluS* upregulated and downregulated genes following *AluJb* overexpression. Gene sets with FDR < 0.05 were considered significant. GSEA: Gene Set Enrichment Analysis, FDR: False Discovery Rate, NES: Normalized Enrichment Score.

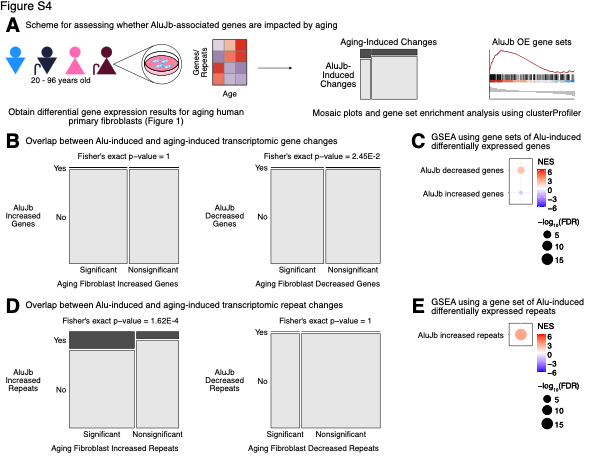

**Supplementary Figure S4. *AluJb*-induced transcriptomic changes are partially induced by aging. (A)** A diagram illustrating how *AluJb*-induced and aging-induced transcriptomic changes were compared. Mosaic plots showing the fraction of *AluJb*-induced (**B)** genes and **(D)** repeats found among significant (FDR < 0.05) and non-significant primary fibroblast aging-induced genes and repeats. Statistical significance of frequency differences was assessed with Fisher’s exact test, and p < 0.05 was considered significant. GSEA analysis with gene sets for *AluJb* upregulated and downregulated **(C)** genes and **(E)** repeats in human aging primary fibroblasts. Gene sets with FDR < 0.05 were considered significant. GSEA: Gene Set Enrichment Analysis, FDR: False Discovery Rate, NES: Normalized Enrichment Score.

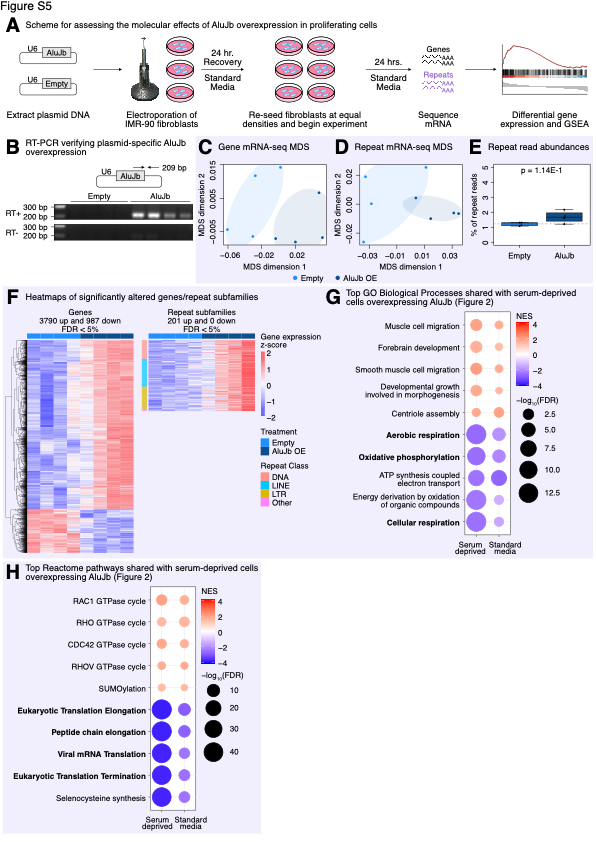

**Supplementary Figure S5. *AluJb* induces similar widespread changes in proliferating and serum-deprived fibroblasts. (A)** A diagram illustrating how the transcriptomic impact of *AluJb* overexpression in proliferating IMR-90 fibroblasts was assessed. **(B)** Plasmid-specific *AluJb* overexpression was assessed by endpoint RT-PCR of empty vector and *AluJb* overexpressing IMR-90 fibroblasts using primers targeting the 3’ *AluJb*-plasmid backbone junction. PCR reactions were carried out with (RT+) or without (RT-) reverse transcription in N = 4 samples per group, and all of these were used for transcriptomic profiling. Multidimensional scaling (MDS) analysis of the **(C)** gene and **(D)** repetitive element transcriptomes across samples. **(E)** Quantification of the percent of total reads mapping to repetitive elements. Statistical significance was assessed with a Wilcoxon rank sum test. **(F)** Gene expression heatmaps for significant (FDR < 0.05) differentially expressed genes and repeat subfamilies. The top 5 significant (FDR < 0.05) and commonly regulated **(G)** GO Biological Process and **(H)** Reactome pathway gene sets in serum-deprived fibroblasts and proliferating fibroblasts in standard media. Fisher’s method was used to combine p-values from GSEA analyses in each media condition, and pathways were ranked on their meta-analysis p-value. RT: Reverse Transcription, GSEA: Gene Set Enrichment Analysis, FDR: False Discovery Rate, NES: Normalized Enrichment Score.

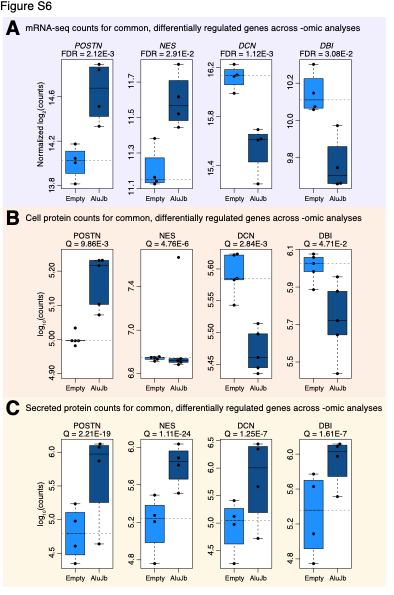

**Supplementary Figure S6. Genes and proteins differentially regulated across “omic” analyses.** The abundances of four significantly altered genes—*POSTN, NES, DCN,* and *DBI*—and their proteins in the **(A)** transcriptome, **(B)** cell proteome, and **(C)** secretome. FDR: False Discovery Rate, Q: Q-value

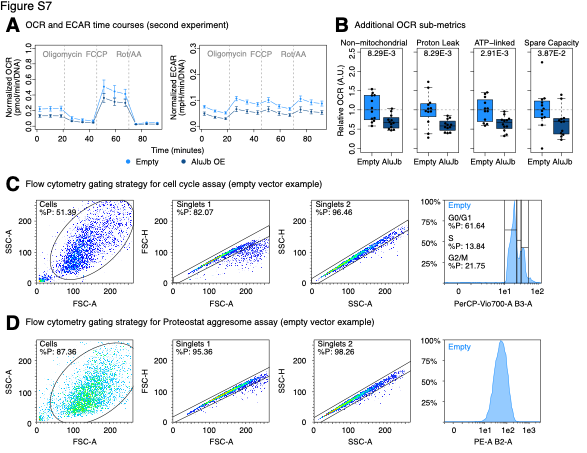

**Supplementary Figure S7. Additional functional profiling data and parameters. (A)** OCR and ECAR time courses from a second experiment with N = 6 independent transfections per group, for a total of N = 12 samples per group. **(B)** Additional OCR sub-metrics, including non-mitochondrial respiration, ATP-linked respiration, proton leak, and spare capacity, in empty control and *AluJb*-overexpressing IMR-90 fibroblasts (N = 12 per group). Statistical significance was assessed with a Wilcoxon rank sum test, and p < 0.05 was considered significant. The flow cytometry gating strategies for the **(C)** propidium iodide-based cell cycle assay and the **(D)** Proteostat aggresome detection assay. OCR: Oxygen Consumption Rate, ECAR: Extracellular Acidification Rate.

**SUPPLEMENTARY TABLE LEGENDS**

**Supplementary Table S1. Transcriptomic analysis of human aging primary fibroblasts.**

**(A)** All DESeq2 results for primary fibroblasts across age using quantifications from TEtranscripts. **(B)** All DESeq2 results for primary fibroblasts across age using quantifications from TElocal. **(C)** All DESeq2 results for primary fibroblasts across endogenous AluJb expression levels using quantifications from TEtranscripts. **(D)** GSEA results (FDR < 0.05) for elevated AluJb expression with GO Biological Process gene sets. **(E)** GSEA results (FDR < 0.05) for elevated AluJb expression with Reactome pathway gene sets.

**Supplementary Table S2. Differential abundance analyses across “omic” layers.**

**(A)** Data-independent acquisition (DIA) isolation scheme. **(B)** Abundances of quantifiable cellular protein groups. **(C)** Abundances of quantifiable secreted protein groups. **(D)** All DESeq2 results comparing the transcriptomes of control and AluJb overexpression groups. **(E)** All differential abundance results between control and AluJb overexpression groups for cellular proteins. **(F)** All differential abundance results between control and AluJb overexpression groups for secreted proteins. **(G)** GSEA results (FDR < 0.05) for AluS-regulated genes in the AluJb transcriptome.

**Supplementary Table S3. GSEA for aging and senescence gene sets across “omic” analyses.**

**(A)** All GSEA results for senescence and aging gene sets in the transcriptome. **(B)** All GSEA results for senescence and aging gene sets in the cellular proteome. **(C)** All GSEA results for senescence and aging gene sets in the secretome. **(D)** All GSEA results for AluJb induced genes and repeats in the human aging primary fibroblast transcriptome.

**Supplementary Table S4. Functional enrichment analyses across “omic” layers.**

**(A)** All significant (FDR < 0.05) GO Biological Process GSEA results in the transcriptome. **(B)** All significant (FDR < 0.05) GO Biological Process GSEA results in the cellular proteome. **(C)** All significant (FDR < 0.05) GO Biological Process GSEA results in the secretome. **(D)** All significant (FDR < 0.05) Reactome pathway GSEA results in the transcriptome. **(E)** All significant (FDR < 0.05) Reactome pathway GSEA results in the secretome. **(F)** All DESeq2 results comparing control and AluJb overexpression groups in proliferating IMR-90 fibroblasts. **(G)** Significant and similarly regulated GO Biological Process GSEA results for serum-deprived and proliferating fibroblasts. **(H)** Significant and similarly regulated Reactome pathway GSEA results for serum-deprived and proliferating fibroblasts. **(I)** All transcription factor regulon enrichment results using decoupleR on the serum-deprived AluJb transcriptome.

**Supplementary Table S5. Multi-contrast gene set enrichment analysis integrating “omic” layers.**

**(A)** All Mitch results with GO Biological Process gene sets. **(B)** All Mitch results with Reactome pathway gene sets.
